# Locus-specific transcriptional regulation of transposable elements by p53

**DOI:** 10.1101/2025.10.01.679811

**Authors:** Julia M. Freewoman, Andrew J. Rosato, Thomas M. Russell, Feng Cui

**Affiliations:** Thomas H. Gosnell School of Life Sciences, Rochester Institute of Technology, 85 Lomb Memorial Drive, Rochester, NY 14623, USA

**Keywords:** p53, transposable elements, locus-specific, transcriptional regulation, chromatin context

## Abstract

The tumor suppressor p53 protects genomic integrity in part by regulating transposable elements (TEs). Most studies of p53-TE interactions rely on synthetic DNA and reporter assays, which estimate expression only at the family or subfamily level and lack locus-specific resolution. To address this limitation, we developed a computational pipeline for ChIP-seq and RNA-seq analysis that employs advanced algorithms to accurately assign short reads mapping to multiple genomic locations. This approach enables precise quantification of TE transcripts at the locus level. By integrating p53 ChIP peaks with differentially expressed TE transcripts, we identified retroelements directly regulated by p53. Applying this framework to lung fibroblast IMR90 and colon cancer HCT116 cells treated with p53 activators, we observed a striking pattern: TEs were predominantly activated in normal IMR90 cells but repressed in HCT116 cancer cells. Analysis of 24 transcriptomes and 10 cistromes further confirmed this trend, distinguishing normal from cancer cell regulation. At the family level, normal cells showed broad TE upregulation, whereas cancer cells exhibited selective repression of Alu and LINE elements, with weaker effects on endogenous retroviruses. These findings provide the first comprehensive, locus-specific view of p53-mediated TE regulation, highlighting distinct chromatin-dependent dynamics in normal and cancer cells.

## Introduction

About 50% of the human genome is composed of transposable elements (TEs) (1). Based on the mechanisms of transposition, TEs are assigned to one of two classes (2). Class I TEs, also named retrotransposons, include long interspersed nuclear elements (LINEs), short interspersed nuclear elements (SINEs), and endogenous retrovirus (ERV) with long terminal repeat (LTR). This class of TEs adopts the ‘copy-and-paste’ transposition mechanism, involving transcription to RNA, reverse transcription to DNA, and reintegration into the genome. Class II TEs include DNA transposons, following the ‘cut-and-paste’ transposition mechanism. The ability of TEs to move their positions across the genome poses a threat to genomic integrity, prompting the development of defense strategies to suppress TE mobilization (3, 4). Therefore, in general, TEs are under tight regulation in somatic cells, although under certain cellular conditions, such as germline formation, preimplantation development, and tumorigenesis, specific TE families are highly expressed (5–9). TEs act as a key regulator of gene expression by providing regulatory sequences (10). Various transcription factors, including GATA (11), LyF-1 (11), Sp1 (12, 13), YY1 (13, 14), and retinoic acid receptors (15), have binding sites (BSs) residing in TEs (16).

The tumor suppressor p53, often referred to as the “guardian of the genome”, is a DNA-binding transcription factor (17) whose levels and activity are induced in response to various stresses, including DNA damage (18). It interacts with DNA in a sequence-dependent manner, with a consensus motif comprising 2 decanucleotides RRRCWWGYYY (R = A, G; Y = C, T; W = A, T), separated by a variable spacer (19, 20). Early studies used the reporter-gene method to identify p53 BSs in vitro (20). Later, p53 BSs were mapped by ChIP-seq experiments across the genomes in vivo (21), resulting in at least 25 p53 cistromes across different cell types (22). Additionally, p53 BSs have been predicted computationally based on position weight matrices (23, 24) or *de novo* motif search (25).

The association between p53 BSs and TEs has been revealed in many studies. For instance, many predicted p53 BSs were found in LINE-1 elements, some of which are functional for L1 transcription upon p53 binding (26). Experimental p53 ChIP sites were also found in LTR (27) and SINE (28) family members. Our analysis of 160 functional p53 response elements showed that 15% of these elements are associated with repeats, with SINE/Alu elements being the most common (24). Furthermore, we performed a meta-analysis of 25 p53 cistromes and found that over 40% of high-confidence ChIP sites reside within TEs, illustrating a strong link between p53 and TEs (22).

The presence of p53 BSs within TEs highlights the regulatory potential of p53 in controlling the expression of TEs. One of the earliest pieces of evidence was provided by Leonova and colleagues (29), which revealed the expression regulation of repetitive DNA elements in cancerous cells by p53. They showed that a mutant p53, but not wild-type p53, resulted in a large increase in the transcription of B1/B2 SINEs in cancer cells, suggesting that p53 represses the expression of SINEs. Moreover, p53 plays a repressive role in controlling LINE retrotransposons in various contexts, including in germline tissues of flies and zebrafish, as well as in human somatic cell lines (30). For certain LINE subfamilies such as L1, p53 constitutively represses transcription by binding to their 5′ untranslated regions (UTRs) and promoting the local deposition of repressive histone marks (31). Wild-type p53 has also been found to repress the transcriptional activity of HERV-I LTR elements, while a mutant p53 variant (V143A) stimulates their expression (32). However, in other studies, wild-type p53 enhances the transcription of LINEs and LTR elements (26, 33, 34). Recent studies found that TE-derived transcripts from LINEs and LTR elements can be either activated or repressed by p53 (35). These findings underscore the critical role of p53 in controlling retrotransposon expression, demonstrating its dual function as an activator or a repressor depending on the cellular context and the types of TE elements. This regulation is essential for maintaining genomic stability and protecting cells from the deleterious effects of uncontrolled TE activity.

Current studies often use synthetic TEs and employ traditional techniques such as reporter assays to estimate TE expression at the family or subfamily level (29–31), which are unable to provide genome-wide analysis of TEs with a locus-level resolution. Meanwhile, high-throughput techniques such as ChIP-seq and RNA-seq have proven difficult to characterize TEs because it is hard to define the origin of short reads that align equally well to multiple genomic locations (36). Because these reads originate primarily from repetitive regions, commonly used analysis pipelines often discard the ambiguously mapped reads, thereby excluding critical regulatory events involving TEs. While recent studies have built tools to handle these reads from RNA-seq (37–50) or ChIP-seq (51–56) either at the family or locus level, the tools are rarely used together to probe TE regulation by a specific TF.

In this study, we developed a novel computational pipeline that integrates CSEM (51) and Telescope (47) to analyze ChIP-seq and RNA-seq data from normal cell (NC) lines and cancer cell (CC) lines treated with p53-activating drugs. Both methods, based on Expectation-Maximization algorithms, enable accurate, locus-specific quantification of TE expression. We found that p53 predominantly activates TEs in the normal lung fibroblast line IMR90 but represses them in the colon cancer line HCT116. Analysis of 24 transcriptomes and 10 cistromes from published datasets confirmed this pattern as a distinguishing feature between NC and CC lines, consistent with our previous findings on chromatin states at p53 binding sites. Family-level analyses revealed that all TE families tend to be upregulated in NC lines, whereas in CC lines, SINEs (particularly Alu elements) and LINEs are primarily downregulated. Interestingly, ERV/LTR elements are more likely to be upregulated in both contexts. Based on these findings, we propose a novel framework for p53-mediated transcriptional regulation of TEs in the chromatin context.

## Materials and Methods

### Genomic datasets

The RNA-seq datasets used in this study include experimental data produced in our lab and those from the literature. Specifically, we produced three RNA-seq data from the normal cell line IMR90, treated by 5-FU for 6, 12, and 24 hr to activate p53 molecules (GSE278889). Together, we analyzed 23 RNA-seq datasets associated with p53 activation (Supplementary Tables S1). Out of the 23 datasets, 12 datasets were from normal cell lines, whereas 11 datasets were from cancer cell lines. Ten published p53 cistromes were used in this study, including seven datasets from normal cell lines and three from cancer cell lines (Supplementary Tables S2). In addition, ChIP-seq data for histone marks H3K4me3 and H3K27ac of both GM12878 and K562 cell lines were obtained from the European Nucleotide Archive (Supplementary Table S3).

### Cell culture

IMR90 p53 +/+ cells (GRCF Biorepository & Cell Center of John Hopkins University) were grown in the recommended Eagle’s Minimum Essential Medium (ATTC) with 10% FBS and 1X antibiotic mix. Cells were grown in the media at 37°C with 5% CO2 in 35 mm plates until 70-80% confluence. At 6, 12, and 24 hours prior to RNA extraction, the media was changed and 2 mL of fresh media, along with 5-FU (with a final concentration of 375 µM) or 2µL dimethyl sulphoxide (DMSO) were added to the cells. Non-treatment controls were also set up, along with biological replicates for each treatment and each time point.

### RNA extraction, sequencing, processing, and analysis

RNA was extracted following the protocol of RNeasy Plus Mini Kit (QIAGEN) with QIAshredder for cell lysis and homogenization. The RNA concentration of the samples was determined using NanoDrop (Thermo Scientific). The RNA samples were sent to the University of Rochester Genomics Research Center for sequencing. FASTQ files obtained from URGRC were processed to remove adapters. Short reads with smaller sizes (< 50 bp) were filtered out, and the remaining reads were further cleaned by FASTX-toolkit with a quality score cutoff of 28. The quality of the resulting data was checked by Cutadapt (57) and FastQC.

The short reads in FASTQ files were aligned to the human reference genome (hg19) using Bowtie2 (58). The assignment of short reads to TEs was performed using Telescope (47) with the aligned reads (in BAM format) and a list of TEs in the gene transcript format (GTF) as inputs. Specifically, the locations of all TEs in the human reference genome (hg19) were obtained from RepeatMasker. The downloaded file was converted to a general feature format 3 (GFF3) file that was subsequently converted to the GTF format using a custom R script. This script created a unique name for each TE by combining the original RepeatMasker name with the chromosome number, as well as the start and stop locations of the TE. The unique names of the TEs allow for the transcript quantification of specific TE loci. The GTF file was created by combining all GFF3 data columns, except for the “Phase” column, and including a custom attribute named “Locus” containing unique TE names. The Telescope program selected the Locus attribute by default as the genomic features where reads were assigned.

The DE analysis of TEs was performed by DESeq2 (59) based on the count of short reads assigned to the elements by Telescope (47). Following the tutorial of the Bioconductor package DESeq2, a sample table was created (Supplementary Table S4). All TEs with total counts for all samples less than 5 per million were excluded. The DE analyses were performed for non-treatment controls versus 5-FU samples, as well as for DMSO samples versus 5-FU samples. The elements with the adjusted p-value (p_adj) lower than 0.05 were selected.

### ChIP-Seq Read Alignment, multiread processing, and peak calling

The Bowtie2 read aligner was used to align the raw ChIP-Seq reads to the human reference genome (hg19). Index files were precompiled with the prefix hg19_ind, which was given to Bowtie2 during alignment. Alignment with Bowtie2 was run on each of the cleaned input FASTQ files using the command *bowtie2 -p 35 -x hg19_ind -U {input.fastq} -S {output.sam} -k 100*, where *-k 100* instructs Bowtie2 to identify up to 100 possible alternative mapping sites where a given read could map. The aligned reads from the samples were outputted into SAM format files and then converted to BAM format files.

CSEM (51) was downloaded from the Dewey Lab (University of Wisconsin). Read probability scores were calculated for every read in each sample file using the command *run-csem --sam - p 10 input_name.sam 51 output_name_csem*. The corresponding input samples were used in the calculation. Once probability scores were assigned to each read, multi-reads were sampled, meaning all multireads associated with a specific read were removed and only the multiread with the highest probability score was retained. This new set of uni-reads and the highest scoring multireads was produced using the CSEM data processing command *csem-generate-input --sampling --bed input_name.bam output_name_csem_sampling*.

Peak calling was then performed using the Model-based Analysis of ChIP-Seq 2 (MACS2) software suite (60). MACS2 was used to identify differentially expressed peaks between control and treatment samples. Peaks were called for the samples in the datasets using the command *macs2 callpeak -t treatment_sample_1_csem_sampling.bed treatment_sample_x_csem_sampling.bed -c control_sample_1_csem_sampling.bed control_sample_x_csem_sampling.bed -f BED --outdir /path/to/out/dir/csem_sampling/ -n dataset_name_csem_sampling*. The ChIP peaks that are statistically significant with - log(q_value) less than 0.05 were chosen for subsequent analysis.

### Genomic data visualization

Processed ChIP-seq and RNA-seq data (in BAM files) were read into R via Rsamtools (version 1.32.3). The coverage of the reads was computed by GenomicAlignments (61) and exported as a BigWig file via rtracklayer (62). The BigWig files were then uploaded to CyVerse. The track, hub, and genome files were made following the directions as detailed by UCSC Genome Browser. The hub was uploaded to the Track Hubs interface in the UCSC Genome Browser. The height of each viewing track for our RNA-seq dataset was kept the same, so the difference in expression between different samples is clearly seen. In addition, the “add custom track” interface was used to add ChIP-seq and RNA-seq tracks.

### Class assignment of differentially expressed transposable elements

Human repetitive region positions in the human reference genome (hg19) were downloaded from the UCSC Genome Browser. The repeat elements were identified using RepeatMasker and Repbase. The major types of repeat elements were selected for analysis, including SINE (Alu, MIR), LINE (CR1, L1, L2 and RTE), LTR (ERV1, ERVK, ERVL and Gypsy), Simple Repeat ((TG)_n_, (TCG)_n_, (CACTC)_n_, (GAGTG)_n_, and (TATATG)_n_), Low Complexity (C-rich, GC-rich, GA-rich, CT-rich), DNA (MuDR, PiggyBac, TcMar-Mariner, hAT-Charlie). The remaining repeat types were included into the “Other” category.

### Monte Carlo simulation

A Monte Carlo simulation was performed to assess the background level of overlapping ChIP fragments obtained from published studies (Supplementary Table S2) following our previous studies (22). In the simulation, 1,134,047 genomic DNA segments (500 bp in length on average) were randomly selected from the human genome assembly hg38, and the number of fragments overlapped with others was determined. This process was repeated 100 times to compute the percentage of randomly selected DNA fragments that overlapped.

## Results

### Co-localization of p53 genomic binding sites and retrotransposon expression

To identify the TEs regulated by p53 at the locus level, we developed a novel pipeline for analyzing ChIP-seq and RNA-seq data to identify p53 ChIP peaks near the TEs and quantify TE expression, respectively (Figure 1). For the ChIP-seq analysis, we utilized CSEM (51), which considers short reads that map to multiple genomic locations (multi-reads). CSEM improves sequencing depth by assigning fractional counts to multi-reads using a weighted alignment scheme, enabling the identification of novel peaks that would otherwise be missed when relying solely on uniquely mapped reads (uni-reads). The updated counts of short reads were sent to ChIP-seq peak caller MACS2 (60) to find the differential peaks between 5-FU-treated samples and controls. As a result, we identified 144,593 (78%) and 39,412 (20%) peaks in IMR90 and HCT116 cells, respectively, with 2,343 (1%) peaks shared between the two cell lines (Supplementary Figure S1). The higher number of p53 peaks found in IMR90 cells compared to HCT116 cells is consistent with a general observation on NC and CC lines (22), in which p53 peaks found in NC lines are higher than those found in CC lines.

**Figure 1.**
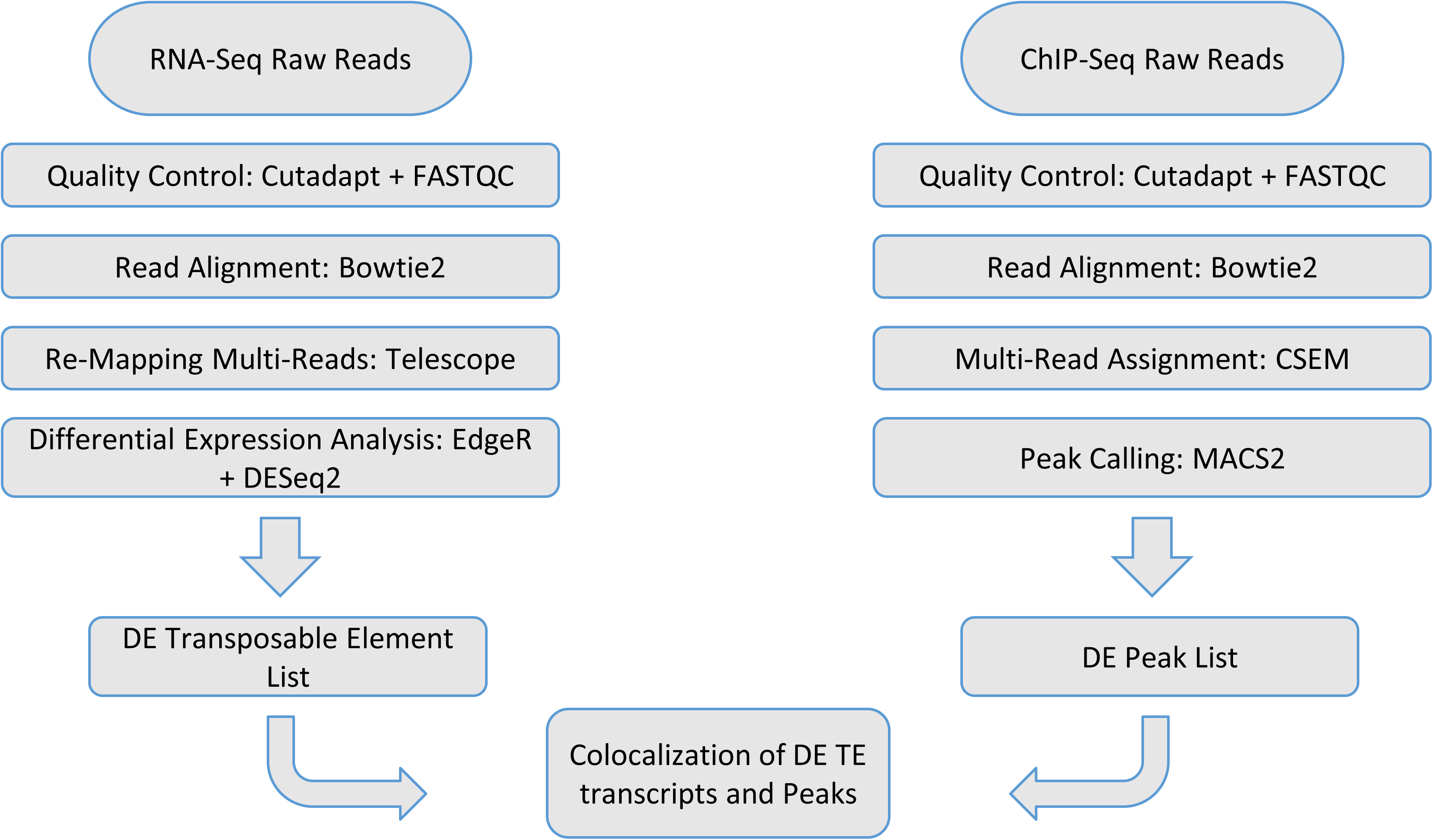
RNA-seq and ChIP-Seq workflows for locus-specific TE quantization. Raw reads were first processed by Cutadapt and FASTQC to remove reads with poor quality. Processed reads were aligned to the genome with Bowtie2. For RNA-seq, multi-reads were realigned to the genome using Telescope. Differentially expressed (DE) transcripts were identified by DESeq2. Resulting DE transcripts were intersected with human TEs from RepeatMasker to identify DE retrotransposons. For ChIP-seq, multi-reads were assigned the most probable locations using CSEM, which, together with uni-reads, were used to call p53 ChIP peaks using MACS2. The differential p53 peaks were colocalized with DE retrotranspons based on their genomic locations. Only the elements that were located within 2 kb from the p53 peaks were chosen, which were considered as p53-regulated TEs.

For RNA-seq data, we first performed quality control using Cutadapt (57) and FastQC to filter out reads with poor quality, which resulted in 30-50 million and 50-80 million high-quality reads for HCT116 and IMR90 cells, respectively (Supplementary Figure S2). Based on alignments from Bowtie2 (58), about 10%, 60% and 30% of reads are unmapped, uniquely mapped (i.e., uni-reads), and ambiguously mapped (i.e., multi-reads), respectively (Supplementary Figure S3). Then we employed Telescope (47), which reassigns multi-reads to the most probable source transcripts as determined within a Bayesian statistical model, to accurately estimate TE expression. DESeq2 (59) was used to identify differentially expressed (DE) transcripts. These transcripts were then intersected with all human TEs from RepeatMasker to identify DE repetitive elements. As a result, we identified more than 11,000 DE elements across all cell lines (Supplementary Table S5). Normal and cancer cell lines have on average, 1,629 and 2,073 such elements, respectively (Supplementary Tables S6 and S7).

Lastly, we co-localized differential p53 ChIP peaks and DE elements to identify p53-associated TEs using GenomicAlignments (61) by using the following criteria: (1) The DE elements should occur in at least two cell lines to reduce false positives; (2) The elements should be within 2 kb from p53 peaks identified by the ChIP-seq pipeline (Figure 1); (3) To simplify the peak-TE relationship, only those elements associated with a single p53 peak are selected (denoted as the “unique” TE set). For the sake of comparison, in the study on HCT116 and IMR90 cells, we also collected the TEs associated with one or more peaks (denoted as the “all” TE set). As a result, we identified 495 and 534 p53-associated TEs in normal and cancer cell lines, respectively (see details below).

### Temporal expression of p53-associated TEs in HCT116 and IMR90 cells

We first determined at which time point the p53-associated TE expression is detectable. We treated IMR90 and HCT116 cells with 5-FU for 6 hours (hr6), 12 hours (hr12), and 24 hours (hr24). Cells treated with DMSO treatment at the same time points are used, along with cells without any treatment. The time-course RNA-seq data of IMR90 and HCT116 cells were analyzed using the pipeline described above (Figure 1). As a result, we identified 195 and 55 TEs differentially expressed in IMR90 and HCT116 cells, respectively (Supplementary Table S8 and S9), which are within 2 kb from p53 ChIP peaks.

Visual inspection of the expression levels of these TEs results in several observations. First, in cells treated with DMSO, TE expression levels remain at similar levels over three time points – hr6, hr12, and hr24, indicating that DMSO treatment leads to no significant changes in TE expression over time (Figure 2A and 2D). Second, in cells treated with 5-FU, the overall TE expression levels are almost the same over the three time points (Figure 2A and 2D). This is because the identified TEs can be either up- or down-regulated, and therefore, the changes in their expression levels are cancelled. Third, up-regulated TEs exhibit a clear upward trend in 5-FU-treated cells over time compared to their counterparts in control samples or DMSO samples (Figure 2B and 2E). Lastly, down-regulated TEs have decreased expression levels in 5-FU-treated cells over time compared with those in control samples and DMSO samples (Figure 2C and 2F).

**Figure 2.**
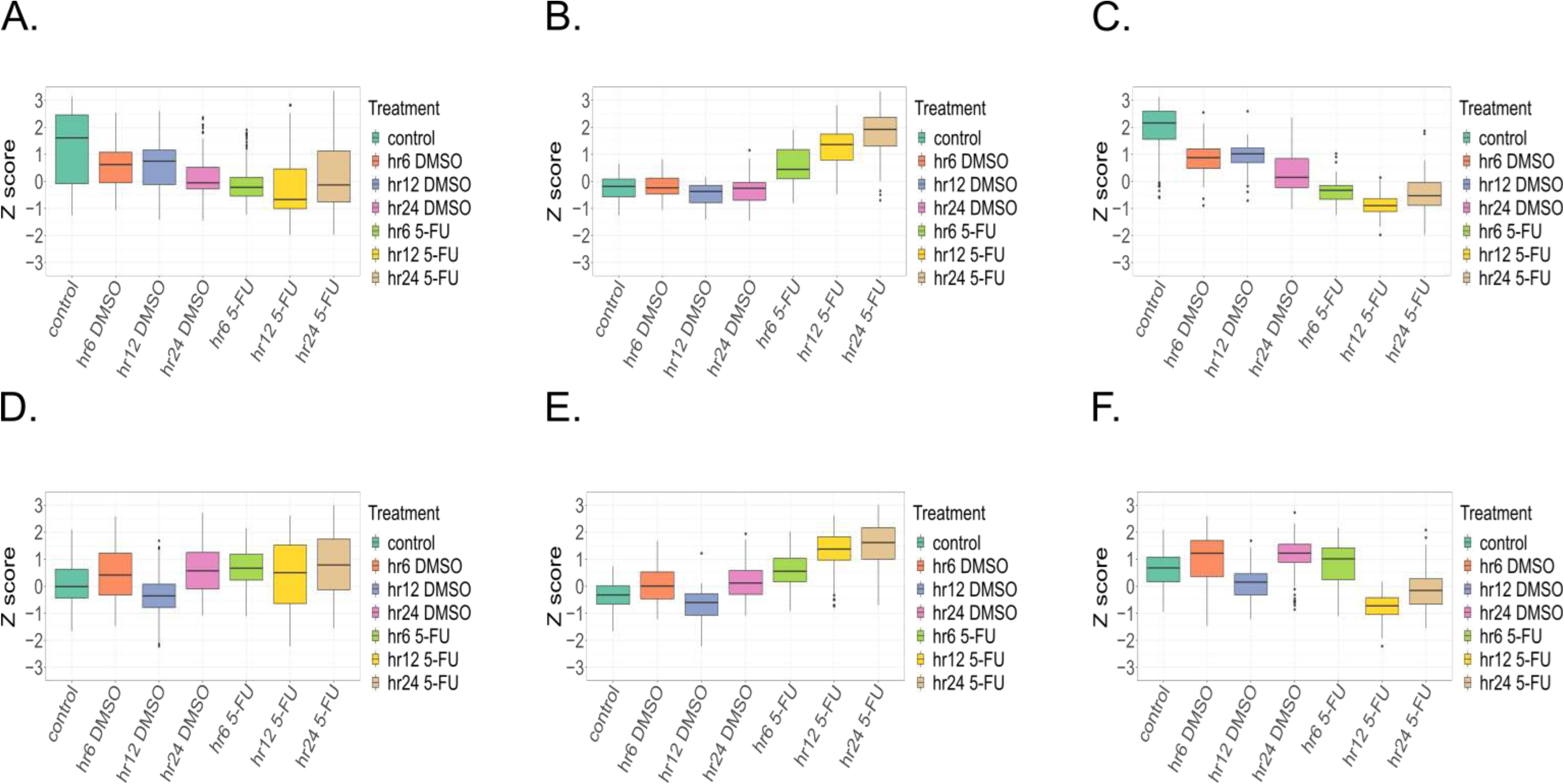
Temporal p53-regulated cell-specific TE expression in HCT116 (A-C) and IMR90 (D-F) cells. TE expression was analyzed for all (A, D), up-regulated (B, E) and down-regulated (C, F) TEs at three time points, 6hr, 12hr and 24hr after 5-FU treatment. Cells with or without DMSO treatment were used as controls. Expression levels of the TEs were obtained by converting read count to Z scores that were subsequently used for comparison. The Z-score values were shown in box-and-whisker plots to indicate median as well as lower and upper quartiles. Only differentially expressed transcripts specific to IMR90 or HCT116 cells were used for analysis and those shared by both HCT116 and IMR90 were excluded.

These observations lead to two conclusions. First, the expression changes of identified TEs over time are associated with the 5-FU treatment, as those changes are not prominent in cells treated by DMSO. Second, detectable changes in TE expression occur at hr12 in both HCT116 and IMR90 cells. Thus, TE expression values at hr12 are used in the subsequent analysis.

### Detailed analysis of TE expression in HCT116 and IMR90

Our previous studies revealed p53 BSs are associated with transcriptionally active histone marks in NC lines, but with repressive histone marks in CC lines (22). We therefore hypothesized that the expression patterns of TEs differ between HCT116 and IMR90 cells. To test this hypothesis, we performed a detailed analysis of cell type-specific TE expressions associated with p53 binding.

Visual inspection was performed on representative elements specifically expressed in HCT116 and IMR90 cells. For example, the AluY_3_305 element downstream of the TRIM27 gene is significantly induced by 5-FU in IMR90 cells (see the rectangle in Figure 3A). The differential expression of this element is clear in the time-course expression plots (Figure 3B-C). That is, in HCT116 cells, the expression levels of this element largely remain constant, except for a bump at hr12, suggesting a transient induction at this time point (Figure 3B). By contrast, in IMR90 cells, its expression levels are steadily increased at hr6 and hr12 and reach a plateau at hr24 (Figure 3C). Conversely, the SVA_D_7_1386 element located upstream of the ZNF586 gene is substantially induced in HCT116 cells (see the rectangle in Supplementary Figure S4A). This element is induced at hr12 and hr24 in HCT116 cells (Supplementary Figure S4B), but not in IMR90 cells (Supplementary Figure S4C). These results clearly showed that TEs exhibit different expression patterns in HCT116 and IMR90 cells upon p53 activation.

**Figure 3.**
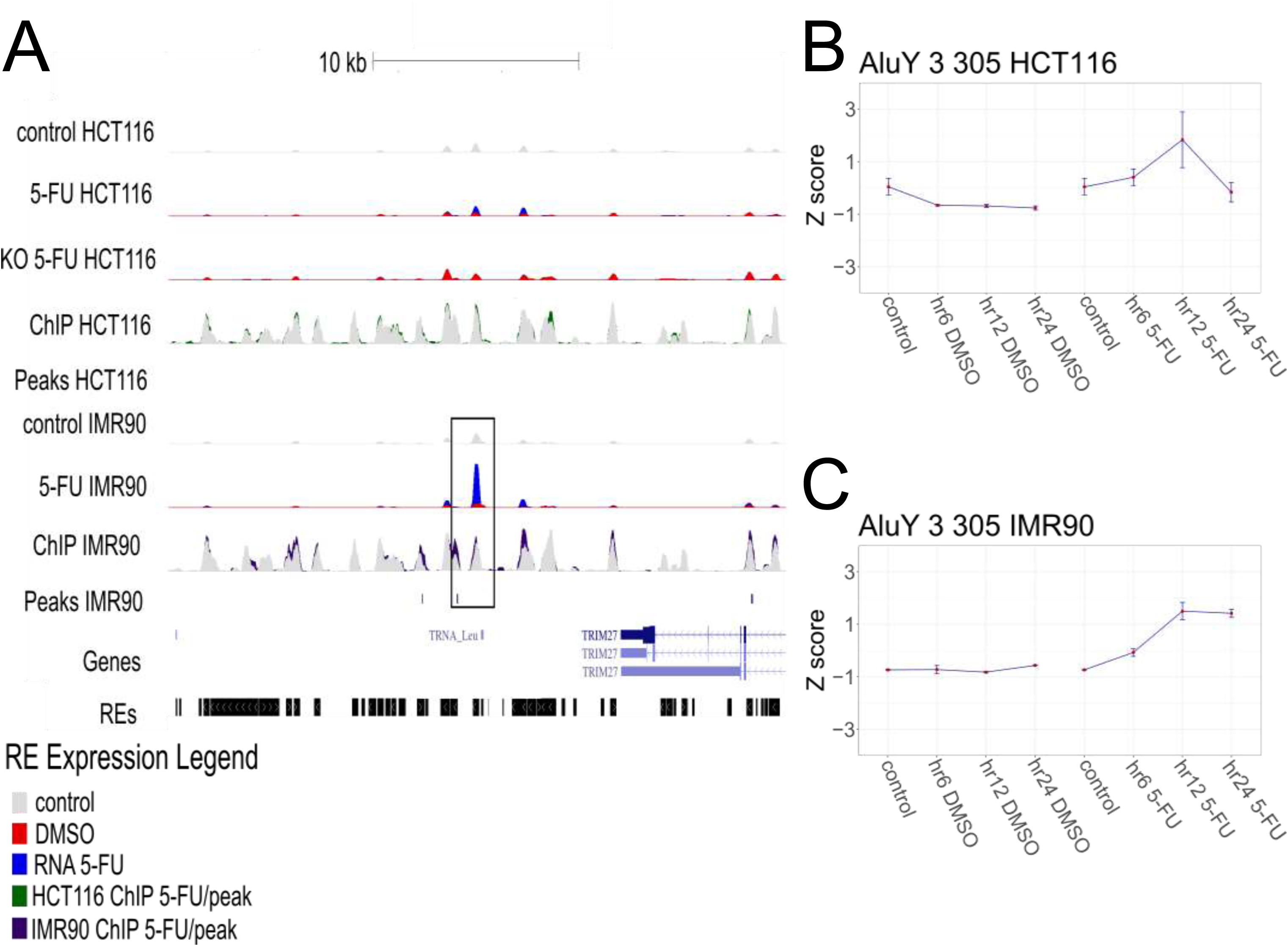
Time-course expression of AluY_3_305 in HCT116 and IMR90 cells. (A) A UCSC Genome Browser view that highlights the time-course TE expression. The session showed the AluY_3_305 locus downstream of the TRIM27 gene. Five tracks displayed at the top show RNA-seq and ChIP-seq datasets from HCT116 cells, followed by four tracks for the corresponding data from IMR90 cells. For RNA-seq data, only the expression values from the time point hr12 were used. The HCT116 datasets included three RNA-seq tracks: control HCT116 with no drug treatment, 5-FU HCT116, and KO 5-FU HCT116, where the HCT116 cells with the TP53 gene knocked-out (KO) were treated with 5-FU. In addition, HCT116 datasets have two ChIP-seq tracks: ChIP HCT116 and Peaks HCT116, which showed read count and peaks called by MACS2, respectively. The IMR90 datasets have two RNA-seq tracks, control IMR90 and 5-FU IMR90, two ChIP-seq tracks, ChIP IMR90 and Peaks IMR90. The tracks showed the average expression level of two biological replicates. The control tracks were shown in gray. For RNA-seq tracks, the DMSO (in red) and 5-FU (in blue) data were superimposed, whereas for ChIP-seq tracks, the DMSO (in grey) and 5-FU (in green for HCT116 or blue for IMR90) data were superimposed. The box on the IMR90 tracks showed the increased expression AluY_3_305 in IMR90 cells, associated with a p53 ChIP peak within 2 kb. There are two other tracks on the browser, one for the UCSC gene track and the other for the repetitive element (RE) track. (B and C) Time-course expression levels of AluY_3_305 in HCT116 (B) and IMR90 (C). The Z scores of the expression values were used to show the normalized expression at hr6, hr12, and hr24 after DMSO and 5-FU treatment, as well as the non-treatment control.

To further characterize these differences at the cell level, we aggregated the up- and down-regulated TEs in the two cell lines. We found that IMR90 cells have more up-regulated TEs (i.e., 107 TEs) than down-regulated ones (i.e., 88 TEs). However, this trend is reversed for HCT116 cells with fewer up-regulated TEs (i.e., 13 TEs) than down-regulated ones (i.e., 42 TEs). This difference is statistically significant with p-value less than 8.7 x 10^-5^ in Chi-square test (Figure 4A, Supplementary Figure S5) and clearly visible in heatmaps (Figure 4B-C, Supplementary Tables S10 and S11).

**Figure 4.**
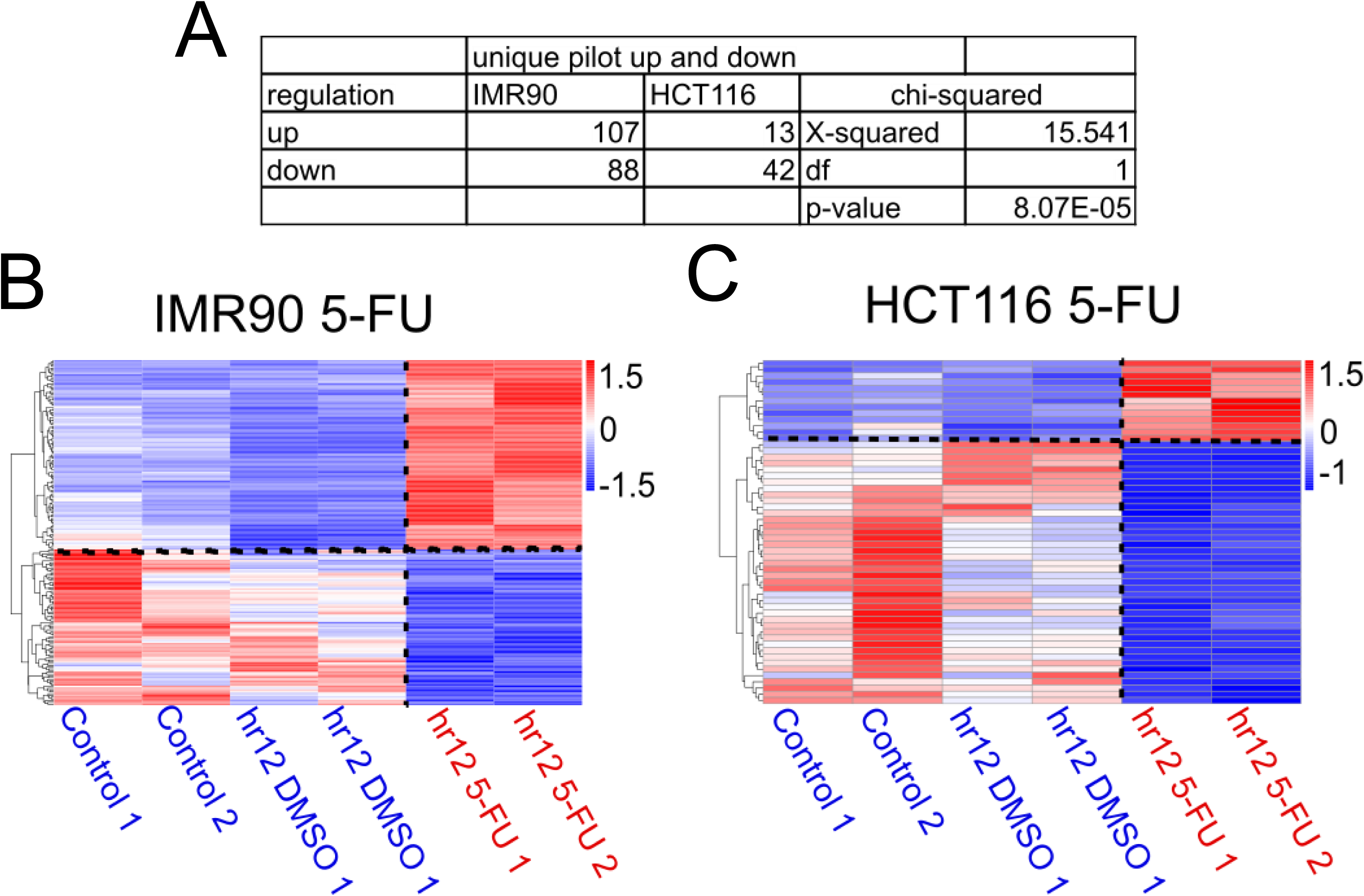
Comparison of p53-associated cell-specific TEs in IMR90 and HCT116 cells. (A) Summary of activated and repressed TEs in IMR90 and HCT116 cells. Statistical significance of the difference between the two cell lines was evaluated by Chi-square test. (B, C) Heatmaps for IMR90 (B) and HCT116 (C). Heatmaps were constructed to compare IMR90 and HCT116 datasets with a dendrogram on the Y-axis to cluster the DE retroelements. Dotted lines indicate the separation between the control (DMSO, blue) and treatment (5-FU, red) samples. Expression levels of the TEs are color-coded.

Note that the “unique” TE sets, in which the TEs are associated with a single p53 ChIP peak, were used in the analysis above. To exclude the possibility that the observed difference only exists for the “unique” TE sets, we repeated the analysis for the “all” TE sets, in which the TEs are associated with one or more p53 peaks. We found the same tendency for both IMR90 and HCT116 cells (Supplementary Figures S6 and S7). Specifically, IMR90 cells have more up-regulated TEs (i.e., 195 TEs) than down-regulated counterparts (i.e., 171 TEs), whereas HCT116 cells have fewer up-regulated TEs (i.e., 55 TEs) than those down-regulated (i.e., 117 TEs) (Supplementary Figure S6A). This difference is also statistically significant (p < 6.0 x 10^-6^, Chi-square test, Supplementary Table 12) and clearly visible in heatmaps (Supplementary Figure S6B-C) and histograms (Supplementary Figure S7). In addition, we examined the time-course expression of these TEs, which clearly showed an overall upward trend for up-regulated TEs and an overall downward trend for down-regulated TEs (Supplementary Figure S8).

To investigate the chromatin landscape around p53-associated TEs in HCT116 (63) and IMR90 (64) cells, we analyzed two enhancer-associated active histone modifications, H3K4me2 and H3K27ac. Previous studies have shown that the two histone marks undergo a marked increase relative to basal levels following p53 activation (63). This gain in the histone marks is substantially reduced after p53 knockdown, suggesting that the deposition of H3K4me2 and H3K27ac around p53 binding sites is highly dependent on p53 local binding. Consistently, in IMR90 cells, both active histone marks are elevated for up-regulated TEs (Figure 5A and 5C), consistent with our previous findings that p53 BSs are associated with active histone marks in NC lines (22). By contrast, H3K4me2 is not elevated in HCT116 cells (Figure 5B), probably related to the epigenetic dysregulation in cancer chromatin (65). For down-regulated TEs, the active marks are not increased around the sites, compared to neighboring regions (+/-1000 bp), except H3K27ac in IMR90 cells (Figure 5D). These results reveal different chromatin landscapes in p53-associated TEs between IMR90 and HCT116 cells, which is in line with the differential expression of p53-associated TEs between the cell lines (Figure 4). These results suggest that such differences may represent a distinguishing feature between NC and CC lines.

**Figure 5.**
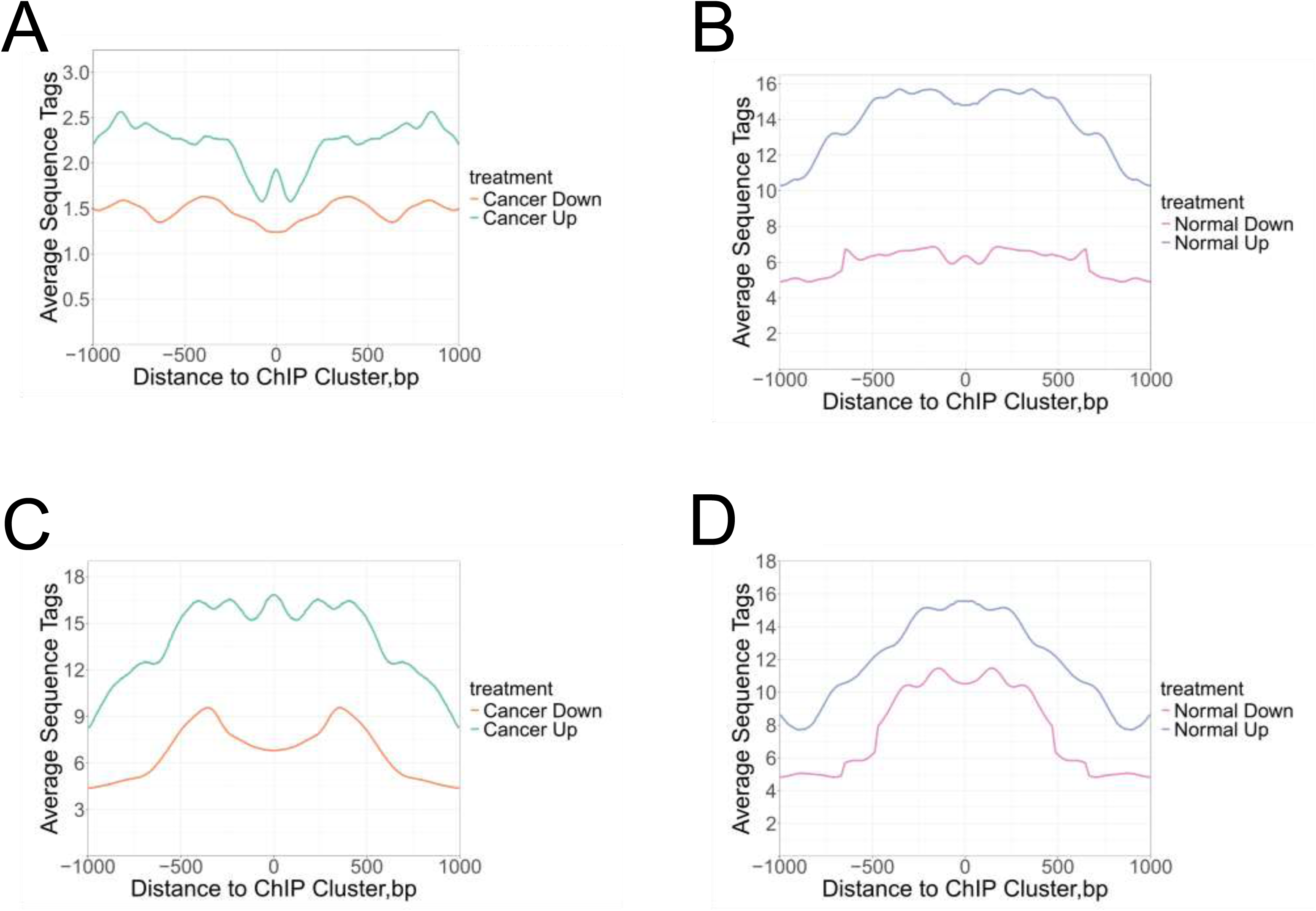
Profiles of the active histone mark H3K4me2 (A, B) and H3K27ac (C, D) around the up- and down-regulated TEs in HCT116 (A, C) and IMR90 (B, D) cells. The aggregate profiles of H3K4me2 and H3K27ac data from HCT116 and IMR90 are shown for the genomic region 1000 bp around the center of p53 ChIP peaks. The averaged H3K4me2 and H3K27ac values were symmetrized across the centers. The ChIP-seq data for histone marks H3K4me3, H3K27me3, and H3K36me3 from both GM12878 and K562 cell lines were downloaded from the European Nucleotide Archive and the UCSC Genome Browser web server. The BAM files of the ChIP-seq data were mapped to human genome assembly hg19. The data were analyzed for four TE groups: normal up, normal down, cancer up, and cancer down (Supplemental Tables S17-S20) following our previous studies (22). The average occupancy values of the histone data were calculated by the dplyr and zoo packages in R. Then, the values were symmetrized with respect to the center of p53 ChIP peaks.

### p53-associated TE expression in normal and cancer cells

To examine this possibility, we performed a comprehensive analysis on 10 cistromes and 23 transcriptomes from multiple NC and CC lines. All cell lines have the wild-type p53 status and were treated with various drugs to activate p53. The pipeline described above (Figure 1) was used to process the ChIP-seq and RNA-seq datasets. The resulting differential p53 ChIP peaks and DE elements were intersected to identify p53-associated TEs.

For the ChIP peaks, we aggregated the peaks from 10 cistromes (Supplementary Table S2) and organized them into clusters, like what we did before (22). The clusters were separated into different bins, denoted as bin-1, bin-2, bin-3, and so forth, based on their coverage, i.e., the number of overlapping members in a cluster (Supplementary Figure S9, Supplementary Table 13). Bin-1 clusters refer to singletons.

We analyzed the height and width of these clusters in the following ways. First, since our clusters contain p53 peaks residing in TEs, we assessed the quality of these clusters by overlapping them with 3,550 and 6,039 consensus p53 ChIP peaks (22) identified in NC and CC lines, respectively (Supplementary Tables S14 and S15). We found that, as the height of the clusters increases, the fraction of overlapping peaks over the total number of peaks in the cluster also increases. For example, bin-5 contains 2,376 peaks, in which 407 peaks (17%) overlap with 3,550 consensus peaks in normal cells (Supplementary Figures S10 and S11). Interestingly, for bin-10, 83 out of 112 (74%) overlap with these consensus peaks. This increase suggests that the peaks in bin-10 are more likely to be found in the consensus sets, highlighting the quality of our data. Second, we calculated the width of the clusters and found that the mean lengths increase from about 100 bp in bin-1 to 800 bp in bin-10 (Supplementary Figure S12), consistent with our observations that bigger clusters tend to be wider. Based on Monte Carlo simulation (Supplementary Table S16), we selected clusters in bin-2 and above as reference p53 ChIP peaks for further analysis.

For the DE retrotransposons, we collated them from 23 cell lines, including 12 NC lines and 11 CC lines (Supplementary Table S1). The elements were co-localized with the reference p53 ChIP peaks, i.e., bin-2 and above, to ensure that they are located within 2 kb of the peaks. To avoid the cell-line-specific DE elements and reduce false positives, we only selected those DE elements that occur in at least two cell lines.

As a result, we identified 491 and 534 DE elements associated with ChIP peaks from NC and CC lines, respectively. We found that NC lines tend to have a higher number of up-regulated TEs (i.e., 284 TEs, Supplementary Table S17) than down-regulated ones (i.e., 204 TEs, Supplementary Table S18). This tendency is reversed for CC lines that have a higher number of down-regulated TEs (i.e., 326 TEs, Supplementary Table S20) than up-regulated ones (i.e., 208 TEs) (Figure 6A, Supplementary Table S19). This difference is also statistically significant (p < 10^-10^, Chi-squared test). Heatmap plots further illustrate the differences (Figure 6B and Supplementary Figure 13, Supplementary Tables S21 and S22). This result is consistent with that from IMR90 and HCT116 cells (Figure 4), suggesting that p53 indeed has a differential impact on TE expression in NC and CC lines.

**Figure 6.**
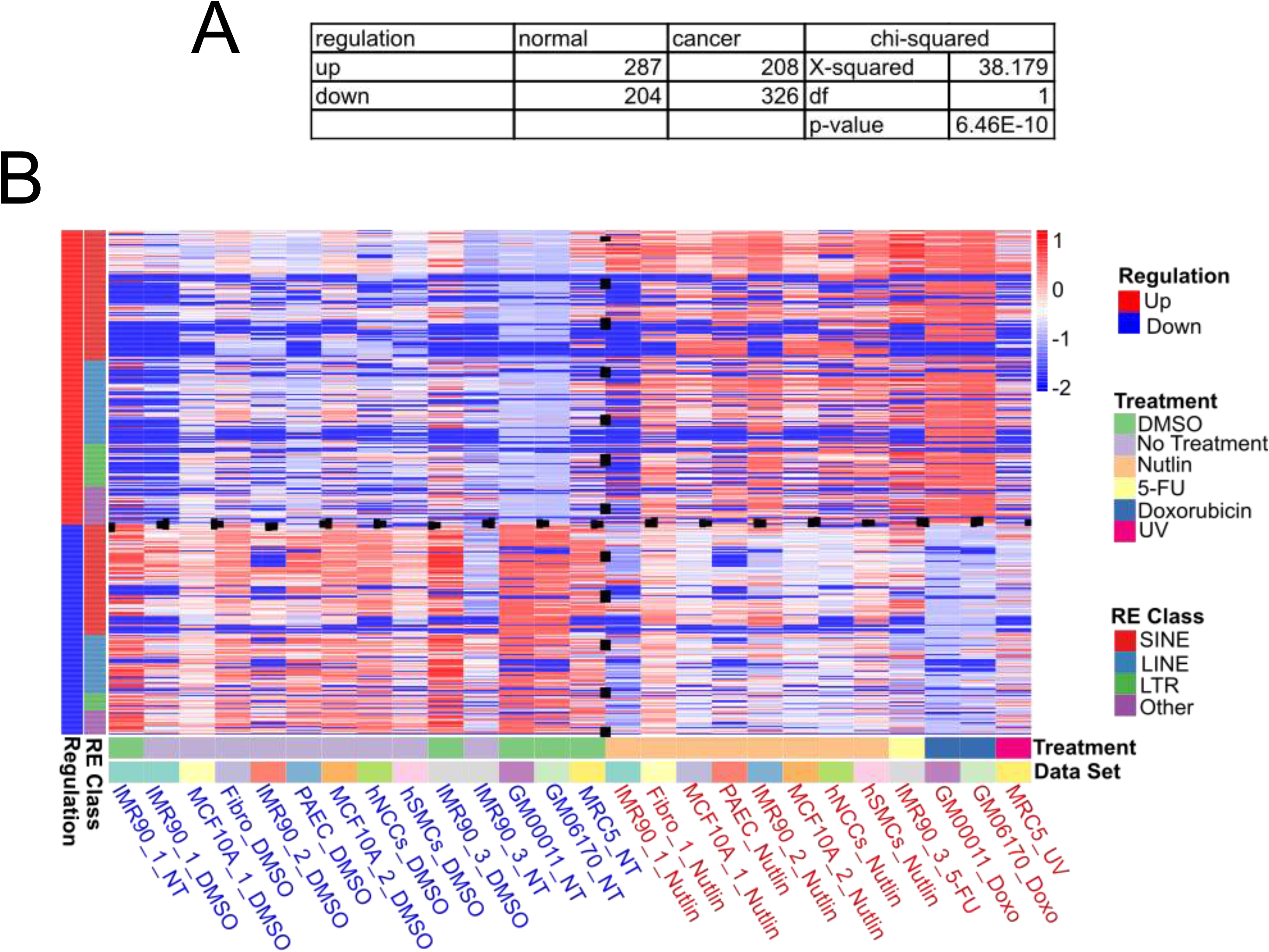
Comparison of p53-associated TEs in normal and cancer cells. (A) Summary of activated and repressed TEs in the NC and CC lines. Statistical significance of the difference between NC and CC lines is evaluated by Chi-squared test. (B) Heatmap for the NC lines. The heatmap is constructed to compare control (DMSO or non-treatment, blue) and treatment (red) samples, as well as TEs up-(red) and down-regulated (blue) by p53. Other notations are the same as those in Figure 4.

To check if the TE differential expression is specific to repeat families, we performed family-level analysis and found that NC and CC lines differ in the family composition in up- and down-regulated TEs (Figure 7). That is, for all repeat families, NC lines have more up-regulated TEs than down-regulated ones. By contrast, CC lines have more down-regulated TEs than up-regulated ones in the SINE and LINE families, but not in LTR and other families (Supplementary Figure S14). These differences are statistically significant (*p* < 0.05, Chi-square test, Supplementary Table S23). Furthermore, we found that Alu elements are highly enriched in down-regulated TEs in the CC lines (Figure 7), among them both AluJ and AluS subfamilies are more pronounced (Supplementary Figure S15). Note that LTR and Other families contain more up-regulated TEs in both NC and CC lines (Figure 7). These results suggest that the regulation of TE expression by p53 is specific to repeat families.

**Figure 7.**
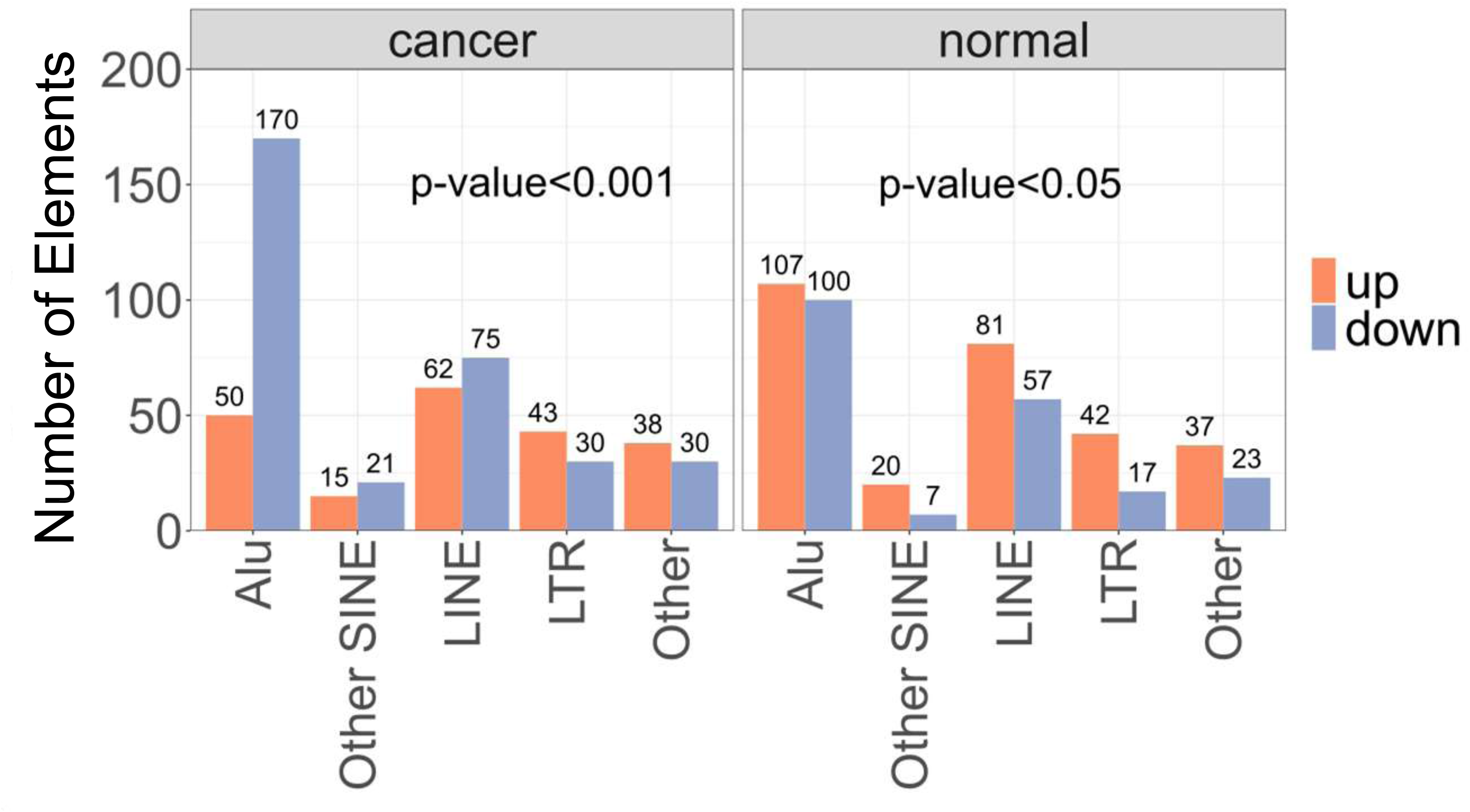
Family-level comparison of p53-associated TEs between normal and cancer cells. Repeat families that the up- and down-regulated TEs in CC (left) and NC (right) lines are counted. For each cell type, the statistical significance of the difference between up- and down-regulated TEs is evaluated by Chi-squared tests.

Lastly, up- and down-regulated TEs differ in the genomic locations in normal and cancer cells (Supplementary Figure S16). Specifically, in normal cells, up-regulated TEs tend to occur in intergenic regions, whereas in cancer cells, down-regulated TEs are predominantly found in UTR regions. The functional significance of such discrepancies requires further investigation.

## Discussion

### A new scheme for locus-specific retrotransposon regulation by p53

Using the computational pipeline we developed to assign multi-reads to TEs (Figure 1), we present the first comprehensive locus-level view of TE regulation by p53. This novel approach enables the identification of individual TEs that are either activated or repressed within repeat families. As a result, we propose a new framework for understanding retrotransposon regulation by p53 at the locus-specific level (Figure 8).

**Figure 8.**
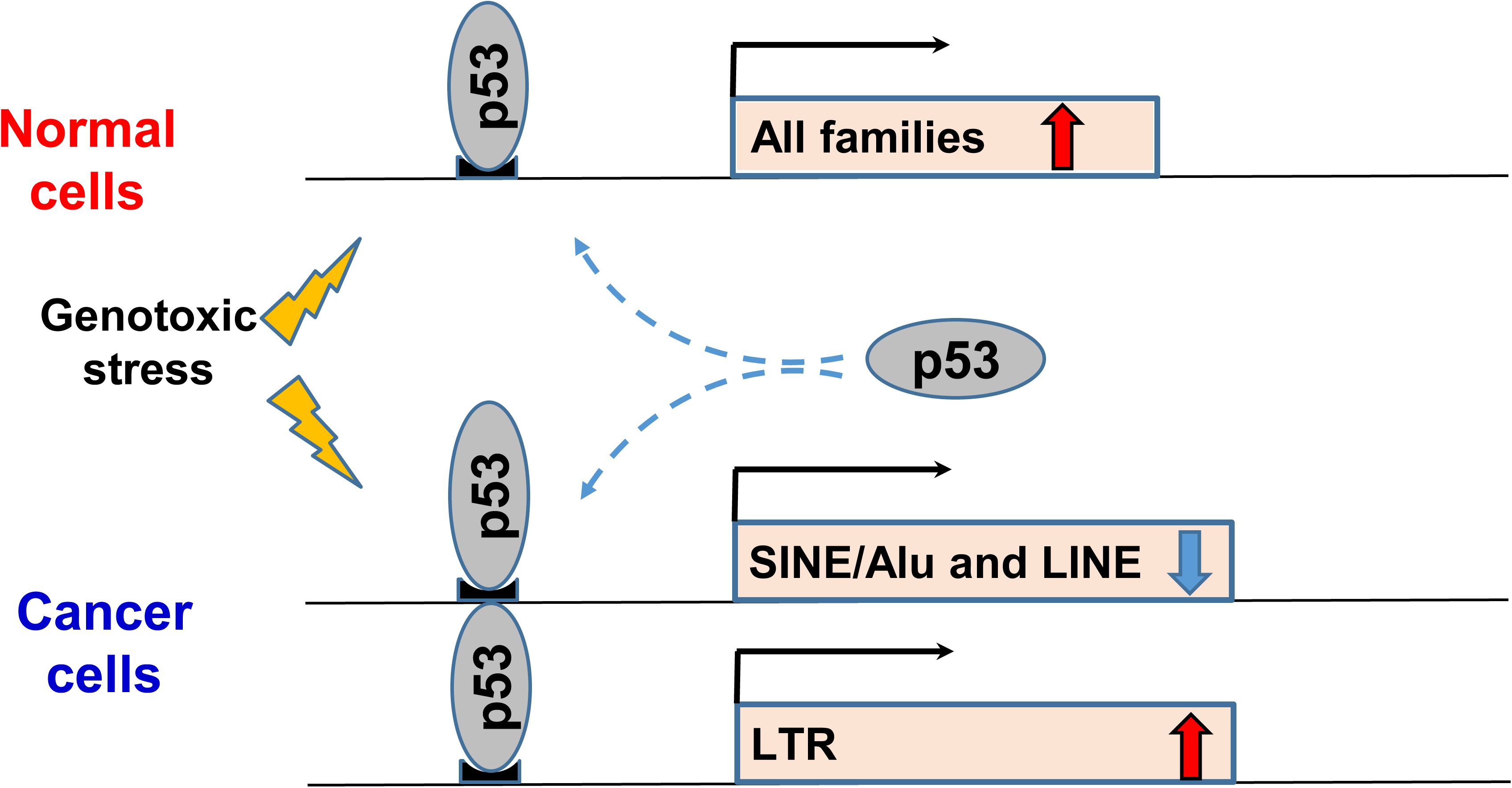
A model for TE regulation by p53 in normal and cancer cells. In normal cells, p53 predominantly activates TEs for all repeat families such as SINE, LINE, ERV/LTR and other families, characterized with active histone marks such as H3K4me2 and H3K27ac. At the level of subfamilies, p53 enhances the transcription of Alu subfamilies such as AluS (Supplementary Figure S15). The preference of activated TEs over repressed TEs occurs in all genomic locations such as UTR, introns and intergenic regions (Supplementary Figure S16). By contrast, in cancer cells, p53 predominantly represses SINEs and LINEs but not ERV/LTR elements. The repression of SINEs is particularly evident for Alu elements (Figure 7), including those in Alu subfamilies such as AluS and AluJ (Supplementary Figure S15). The repressed TEs tend to occur in UTR or intergenic regions (Supplementary Figure S16). Cancer cells are generally characterized by low levels of DNA methylation. Reduction of DNA methylation levels leads to chromatin decondensation, which helps de-repress TE expression (65). Wild-type p53 binds to these elements and represses their expression to antagonize this effect and protect the genome’s integrity. Thus, the shift of TE regulation patterns of p53 in cancer cells from those in normal cells appears to reflect cancer-associated epigenetic dysregulation.

According to our findings, p53 demonstrates distinct family-specific regulatory behavior in NC and CC lines (Figure 7). In NC lines, p53 predominantly activates retrotransposons in response to genotoxic stresses (Figures 4 and 6) across all repeat families (Figure 7). This activation is accompanied by the enrichment of active histone modifications, such as H3K27ac and H3K4me2, around the regulated TEs (Supplementary Figure S16). These results are aligned with our previous studies, which reported that p53 BSs in NC lines are often associated with transcriptionally active histone marks (22).

In CC lines, however, p53 primarily represses elements in specific families, such as SINEs and LINEs (Figure 6), consistent with prior research (29–31). For example, profiles of L1 expression in Wilms tumors showed no expression in tumors with wild-type p53, while tumors with mutant p53 exhibited high L1 expression (30). Similarly, in cancer cell lines like A375, p53 directly represses L1 transcription by binding to their 5’ UTR and promoting the local deposition of repressive histone marks. When p53 or its binding sites in L1 elements are eliminated, these elements are strongly expressed (31). Furthermore, p53 collaborates with DNA methylation and an interferon response to silence B1/B2 SINEs (29). Finally, p53-mediated repression of both SINEs and LINEs has been observed in wild-type p53 CC lines, such as breast cancer (MCF7), osteosarcoma (SJSA-1), and colon cancer (HCT116) (33). Our findings are also in line with the observations that p53 BSs in CC lines tend to reside in genomic regions with repressive histone marks (22). Together, these findings underscore the role of p53-mediated TE suppression as a conserved mechanism of tumor suppression, protecting somatic cells against genomic instability.

One particularly striking observation from our study is the significant repression of Alu elements in cancer cells (Figure 7). The number of repressed Alu elements far exceeds those activated, corroborating our earlier findings (24). We previously identified a hotspot of p53 binding sites near the Pol III promoter of Alu elements (24). Studies have shown that p53 can repress Pol III transcription through interactions between its N-terminal domain and TFIIIB, a factor containing the TATA-binding protein (66–68). Additionally, p53’s DNA-binding domain (DBD) has been shown to be critical for Alu transcriptional repression (69). We proposed a simple mechanism for this repression (24): p53 may bind directly to the hotspot, and consequently, the N-terminal region of p53 would directly interact with TFIIIB, blocking the assembly of the Pol III transcription machinery and thereby silencing Alu transcription. Since most p53 mutations in cancer occur within its DNA-binding domain, the lack of p53 binding leads to an increase of Alu transcription. This mechanism explains the elevated Alu RNA levels and Pol III hypersensitivity observed in many cancers (67, 70).

### Up-regulation of ERV elements by p53 in both normal and cancer cells

ERV/LTR elements, remnants of ancient retroviral infections that occurred millions of years ago, make up approximately 8% of the human genome (71). These elements, particularly newly integrated inserts, are initially subjected to strong DNA methylation, effectively silencing their TF binding sites (72). This methylation-driven repression serves to protect the genome from the potentially harmful effects of ERV elements on gene regulatory networks. However, upon demethylation, some ERV/LTR elements are co-opted as transcribed regulatory elements, expanding the repertoire of functional TF binding sites (73).

Recent studies have revealed that approximately 110,000 ERV elements (about 15%) harbor at least one TF binding site (74), significantly influencing the transcriptional networks of various TFs, including p53. Notably, more than one-third of p53 ChIP-seq binding sites are located within ERVs containing at least one p53 motif (27). Our meta-analysis of 25 published p53 cistromes further demonstrated that ∼40% of p53 ChIP-seq fragments occur within TE families, with a pronounced enrichment in ERV1 elements (22). These findings underscore the significant regulatory potential of p53 on ERV elements.

Early studies on p53-mediated regulation of ERVs focused on the LTR of a human ERV class I element (HERV-1). Reporter assays demonstrated that wild-type p53 inhibits the transcriptional activity of the HERV-1 LTR, whereas a p53 mutant (V143A) stimulates its activity, suggesting a repressive role for p53 in regulating ERV expression (32). Surprisingly, recent studies have shown that p53 activation, triggered by MDM2 inhibitors, can induce the expression of ERVs in cancer cell lines such as MCF-7, SJSA-1, and HCT116 (33). For instance, p53 was found to activate LTR5H elements by directly binding to sites within these elements, and silencing p53 led to reduced transcription (75). Moreover, LTR-derived transcripts can be activated by p53 in cancer cells (35). These findings suggest that activated p53 can either induce or repress the transcription of specific ERV elements in cancer cells, aligning with our data (Figure 7).

While family-level studies on p53-mediated ERV regulation have produced seemingly conflicting results, our locus-level analysis reveals that individual ERV elements can be either upregulated or downregulated by p53. This nuanced perspective provides critical insight into the complex regulatory relationship between p53 and ERVs, further advancing our understanding of how p53 shapes the transcriptional landscape of these retroelements.

## Acknowledgments

The authors would like to acknowledge members of University of Rochester Genomic Research Center for help on library preparation and RNA sequencing, as well as members of RIT Research Computing for computing support.

## Availability of data and materials

All sequencing data for RNA-seq experiments from our lab have been deposited in NCBI’s Gene Expression Omnibus and are accessible through GEO Series accession number GSE157300 and GSE278889.

