## Supplementary Figures for "Locus-specific transcriptional regulation of transposable elements by p53"

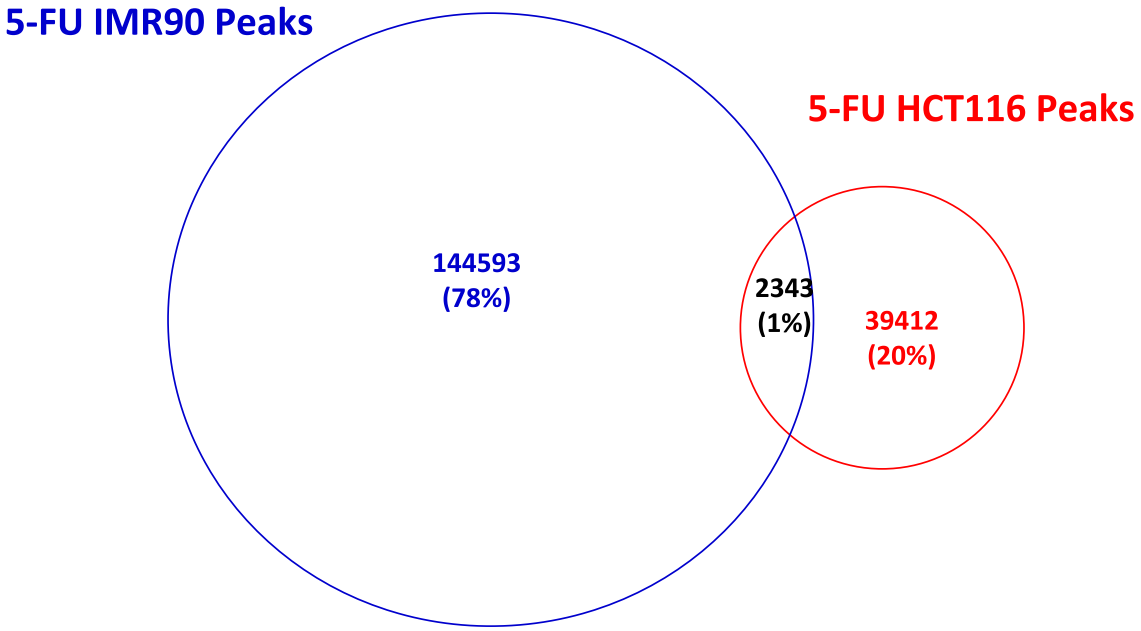

**Figure S1**. Overlap of p53 ChIP peaks between IMR90 cells and HCT116 cells treated with 5-FU. ChIP peaks were called following the procedure described in Figure 1.

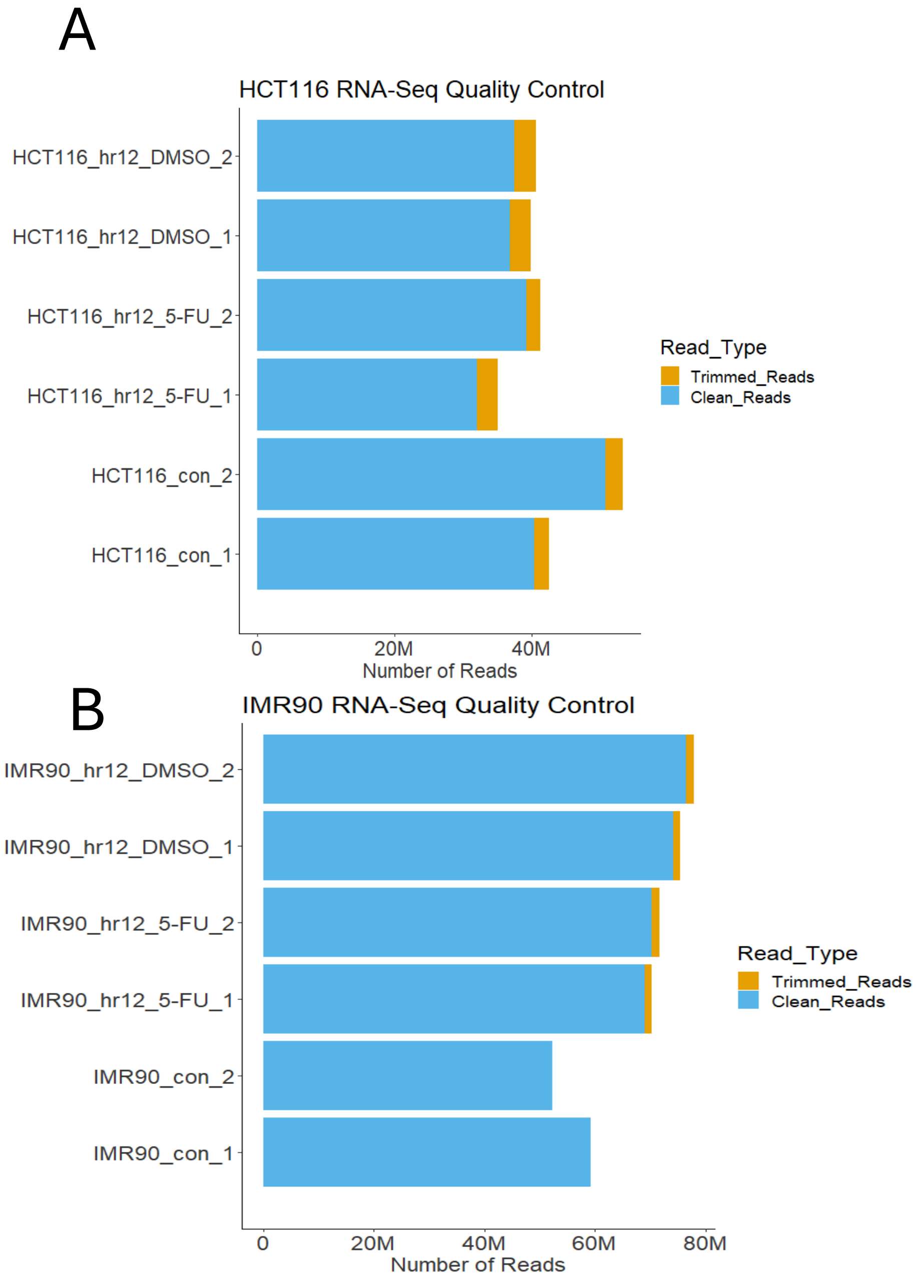

**Figure S2**. Quality control of RNA-seq data from HCT116 (A) and IMR90 (B) cells with 5-FU (12 hr), DMSO, or no treatment. Raw RNA-Seq reads are filtered by the Trimmomatic read quality control software (Bolger et al. 2014) and the amounts of resulting reads filtered-out (yellow) or retained (blue) are demonstrated.

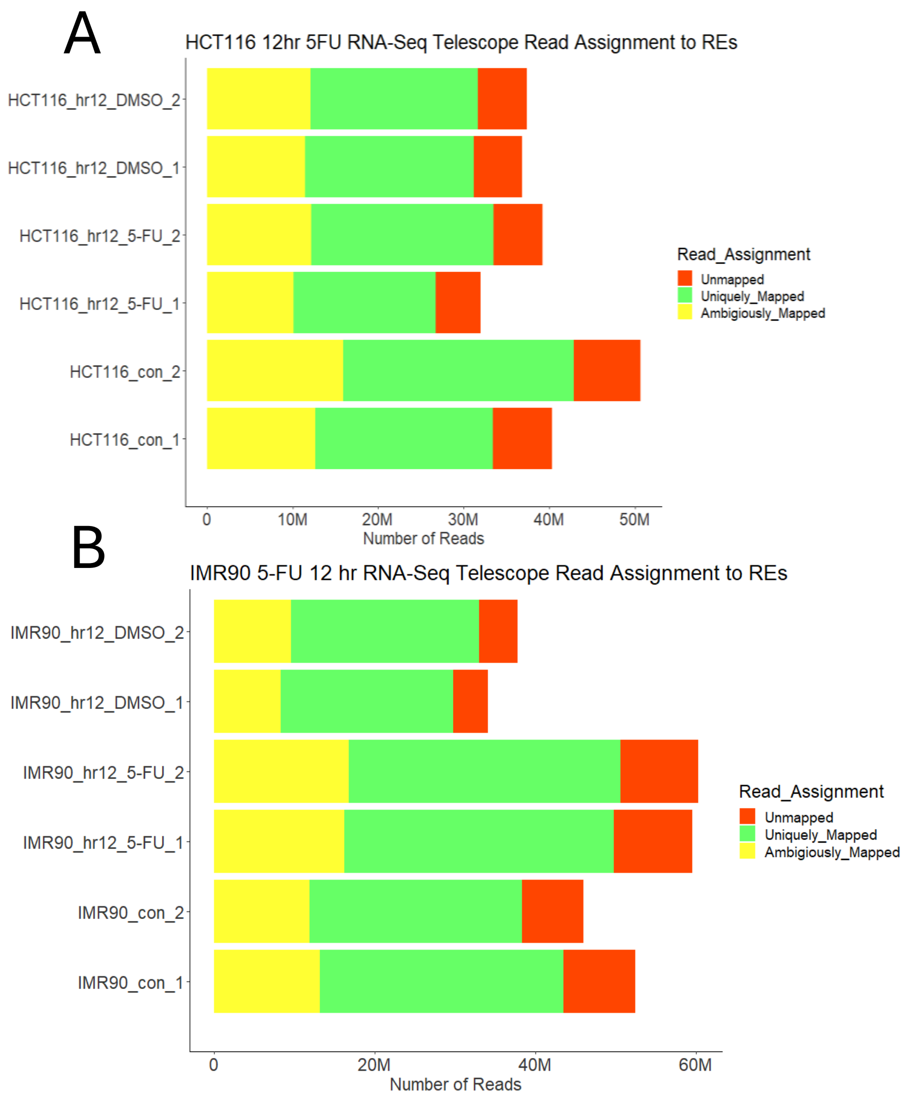

**Figure S3.** Mapping summary of RNA-seq data from HCT116 (A) and IMR90 (B) cells with 5-FU (12 hr), DMSO or no treatment. Bowtie2 is used to map processed RNA-seq reads to the human genome, in which reads are assigned to one of three categories: uniquely mapped positions (green), ambiguously mapped positions (yellow) and unmapped positions (red).

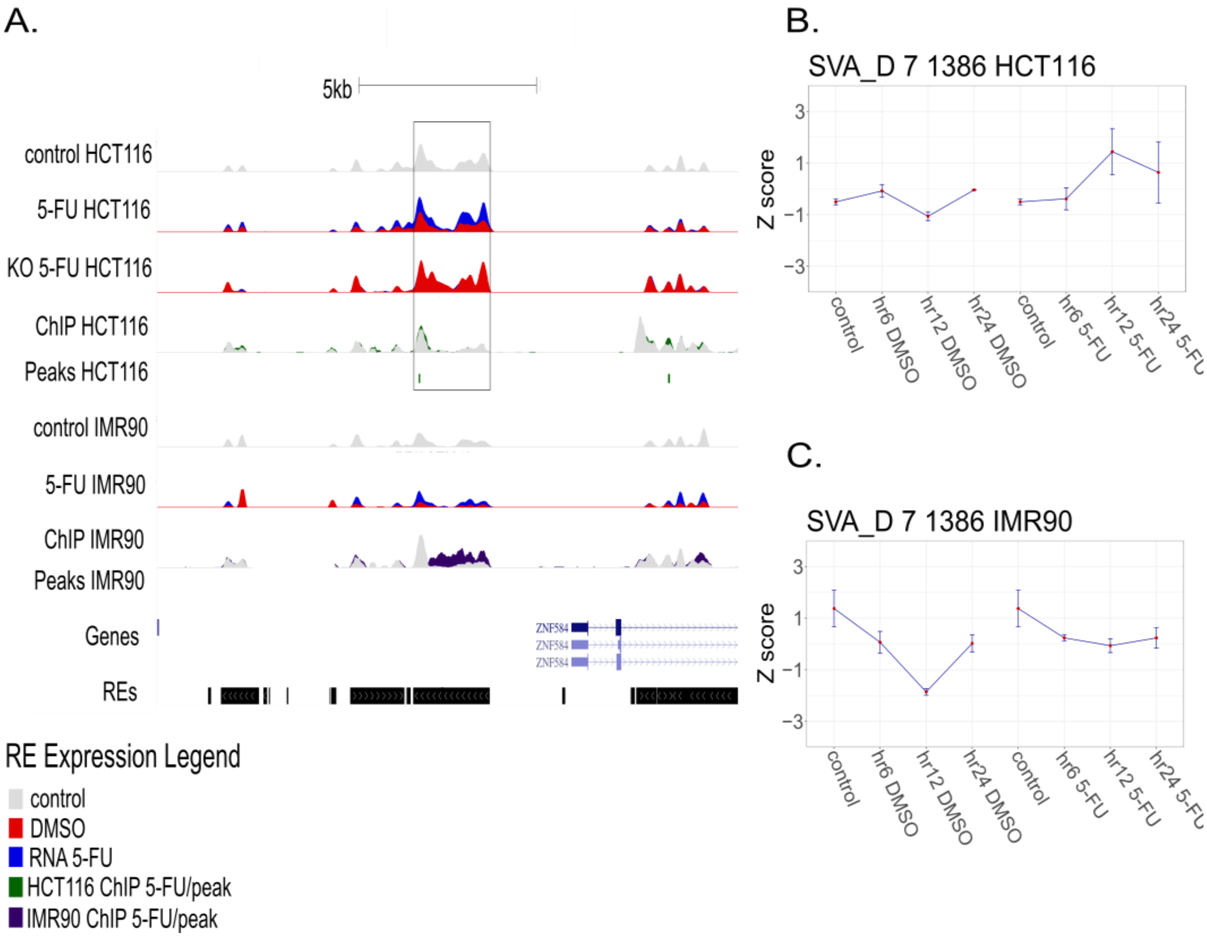

**Figure S4.** Time-course expression of SVA_D_7_1386 in HCT116 and IMR90 cells. (A) A UCSC Genome Browser view that highlights the time-course TE expression. The session shows SVA_D_7_1386 locus downstream of the ZNF584 gene. (B, C). Time-course expression levels of SVA_D_7_1386 in HCT116 (B) and IMR90 (C). Other notations are the same as Figure 3.

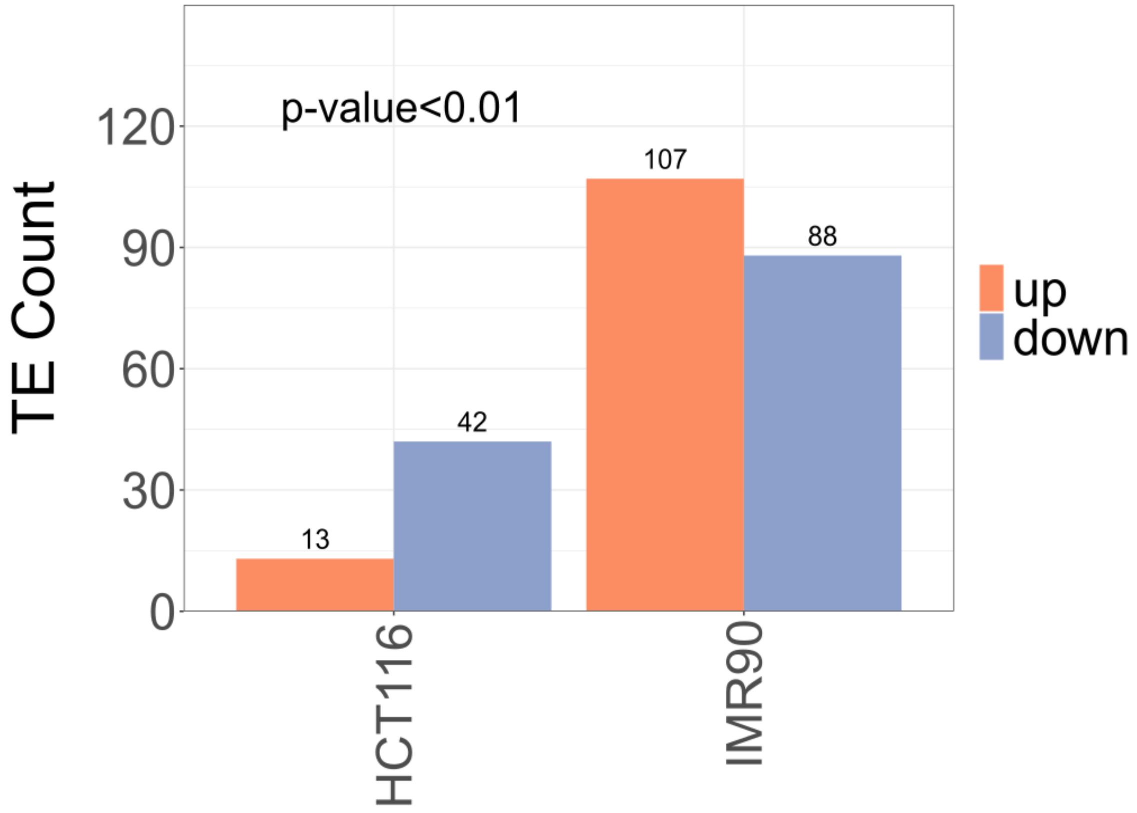

**Figure S5.** Summary of cell-specific TEs regulated by p53 in IMR90 and HCT116 cells. The total number of up- (orange) and down-regulated (blue) TEs in HCT116 cells (left) are compared with that in IMR90 cells (right). The statistical significance of the difference is evaluated by Chi-squared test. Note that TEs analyzed here are associated with one p53 peak.

A.

| | all pilot up and down | | | |
|---|---|---|---|---|
| regulation | IMR90 | HCT116 | chi-squared | |
| up | 195 | 55 | X-squared | 20.497 |
| down | 171 | 117 | df | 1 |
| | | | p-value | 5.97E-06 |

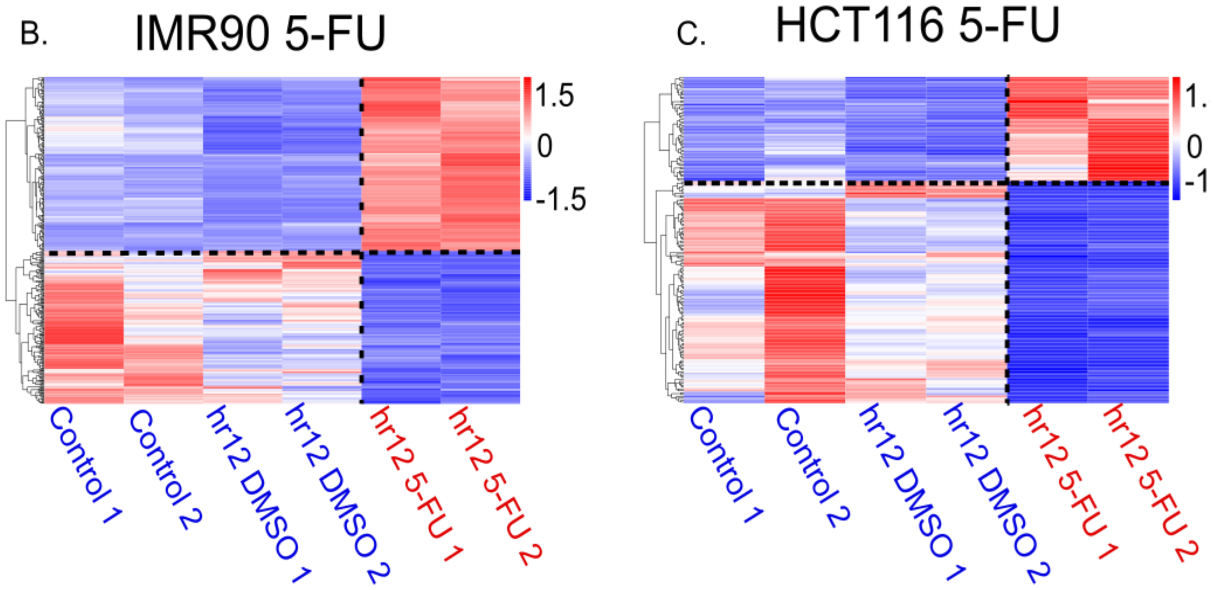

**Figure S6.** Comparison of p53-regulated cell-specific TEs in IMR90 and HCT116 cells. (A) Summary of activated and repressed TEs in IMR90 and HCT116 cells. (B, C) Heatmaps for IMR90 (B) and HCT116 (C). Other notations are the same as Figure 4. Note that TEs analyzed here include the elements that are associated with one or more p53 ChIP peaks.

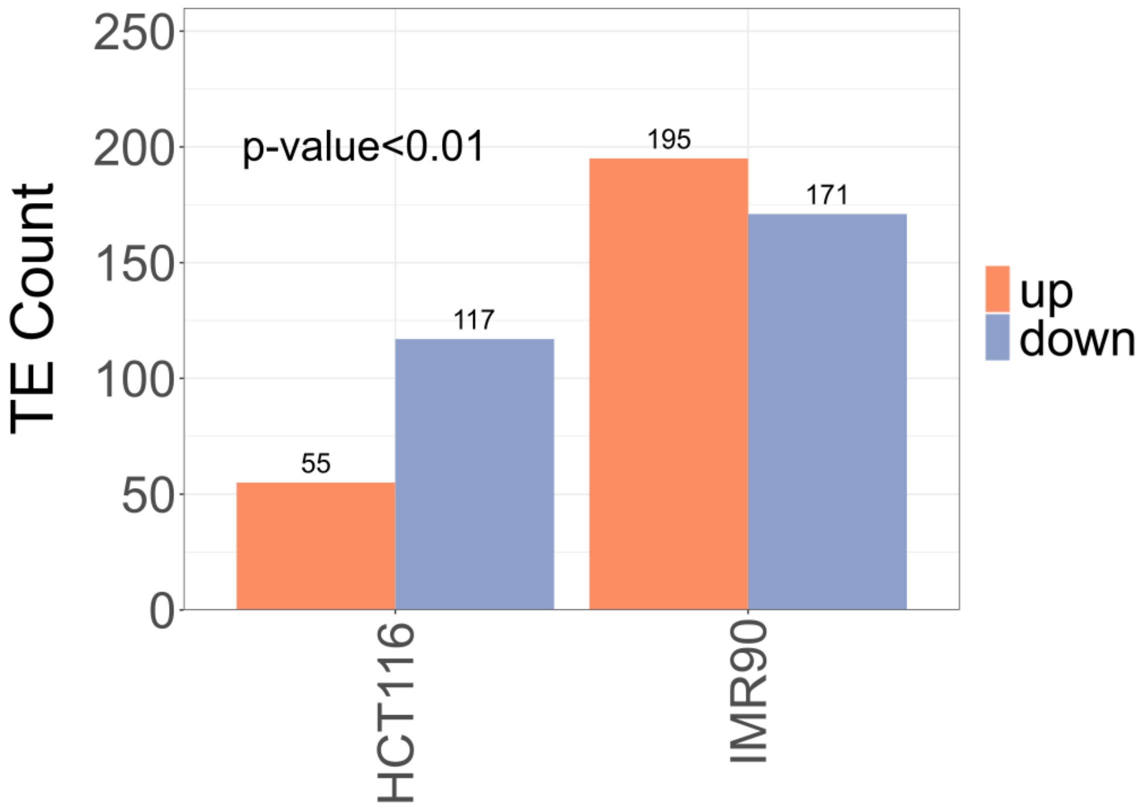

**Figure S7.** Summary of cell-specific TEs regulated by p53 in IMR90 and HCT116 cells. Total number of up- (orange) and down-regulated (blue) TEs in HCT116 cells (left) are compared with that in IMR90 cells (right). The statistical significance of the difference is evaluated by Chi-squared test.

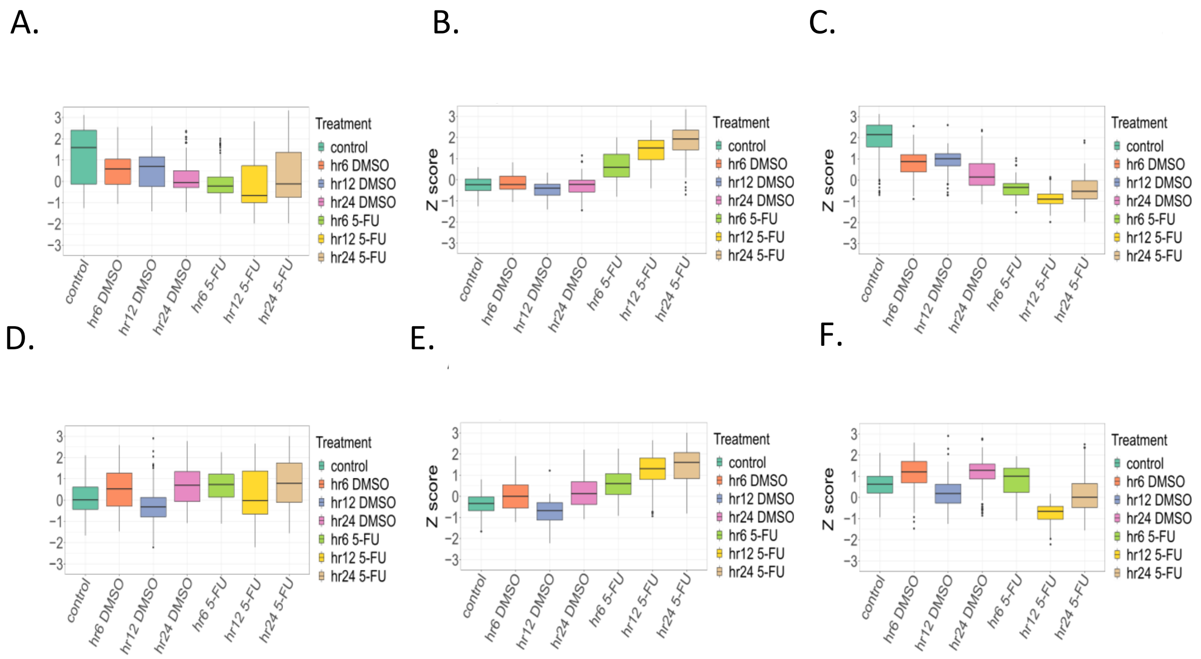

**Figure S8.** Temporal p53-regulated cell-specific TE expression in HCT116 (A-C) and IMR90 (D-F) cells. TE expression is analyzed for all (A, D), up-regulated (B, E), and down-regulated (C, F) TEs at three time points, 6hr, 12hr, and 24hr after 5-FU treatment. Other notations are the same as Figure 2. Note that TEs analyzed here belong to the 'all' set, which includes the elements that are associated with one or more p53 ChIP peaks.

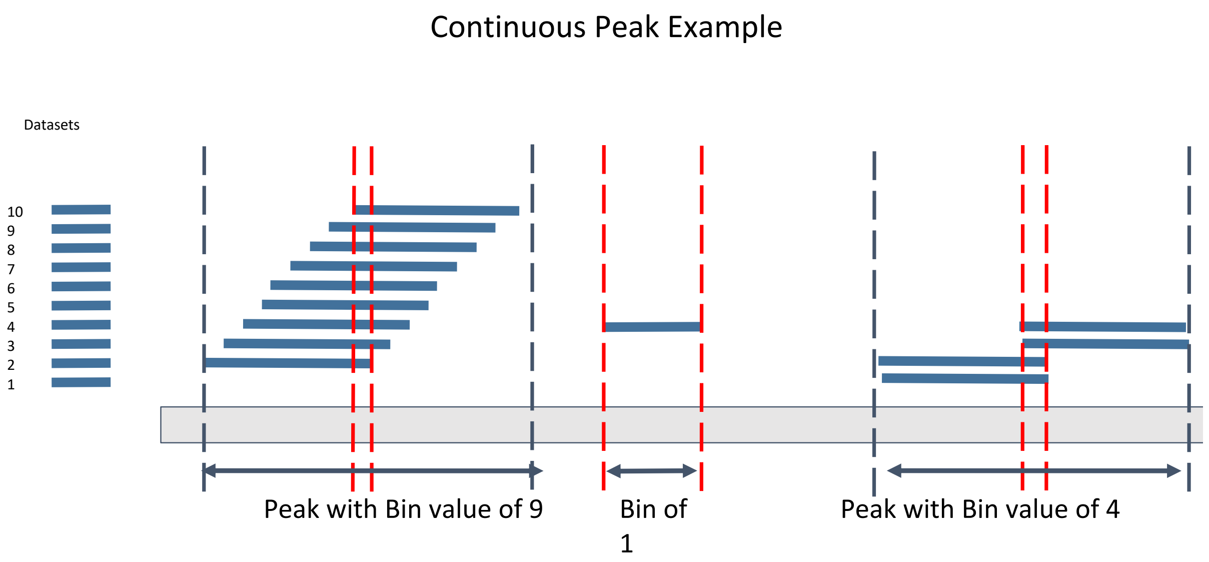

**Figure S9.** Diagram for continuous ChIP peaks. Several examples of continuous peaks are illustrated. Ten ChIP-seq datasets are considered, each with different cell/treatment combinations. Continuous peaks refer to genomic regions that are covered by ChIP peaks from at least one dataset. The value at each position of a continuous peak represents the number of datasets having this position covered by ChIP peaks. All positions in a continuous peak have non-zero values with the maximum value being 10, which is the number of datasets. Each continuous peak is assigned to a bin with the bin value equal to the maximal value of its positions. Red dashed lines show the bin value of the continuous peaks, whereas the black arrows represent the width of the continuous peaks.

A.

| bin_number_vector | intersect_vector | bin_continous_peaks | Bao_total_peaks | pct |
|---|---|---|---|---|
| 1 | 1944 | 661900 | 3550 | 0.002937000 |
| 2 | 702 | 145770 | 3550 | 0.004815806 |
| 3 | 520 | 36813 | 3550 | 0.014125445 |
| 4 | 431 | 8695 | 3550 | 0.049568718 |
| 5 | 407 | 2376 | 3550 | 0.171296296 |
| 6 | 383 | 1057 | 3550 | 0.362346263 |
| 7 | 484 | 1008 | 3550 | 0.480158730 |
| 8 | 388 | 631 | 3550 | 0.614896989 |
| 9 | 321 | 438 | 3550 | 0.732876712 |
| 10 | 83 | 112 | 3550 | 0.741071429 |

B.

| bin_number_vector | intersect_vector | bin_continous_peaks | Bao_total_peaks | pct |
|---|---|---|---|---|
| 1 | 1453 | 661900 | 6039 | 0.002195196 |
| 2 | 732 | 145770 | 6039 | 0.005021609 |
| 3 | 636 | 36813 | 6039 | 0.017276506 |
| 4 | 573 | 8695 | 6039 | 0.065899942 |
| 5 | 556 | 2376 | 6039 | 0.234006734 |
| 6 | 576 | 1057 | 6039 | 0.544938505 |
| 7 | 798 | 1008 | 6039 | 0.791666667 |
| 8 | 568 | 631 | 6039 | 0.900158479 |
| 9 | 426 | 438 | 6039 | 0.972602740 |
| 10 | 107 | 112 | 6039 | 0.955357143 |

**Figure S10.** Overlap between continuous peaks with published consensus ChIP peaks in normal (A) and cancer (B) cells. Continuous peaks in each bin are compared with the normal cell (A) dataset and the cancer cell (B) dataset from previous studies (Bao et al. 2017) and the number of the intersection peaks is shown. The fraction of the intersection peaks over the total number of continuous peaks in each bin is calculated.

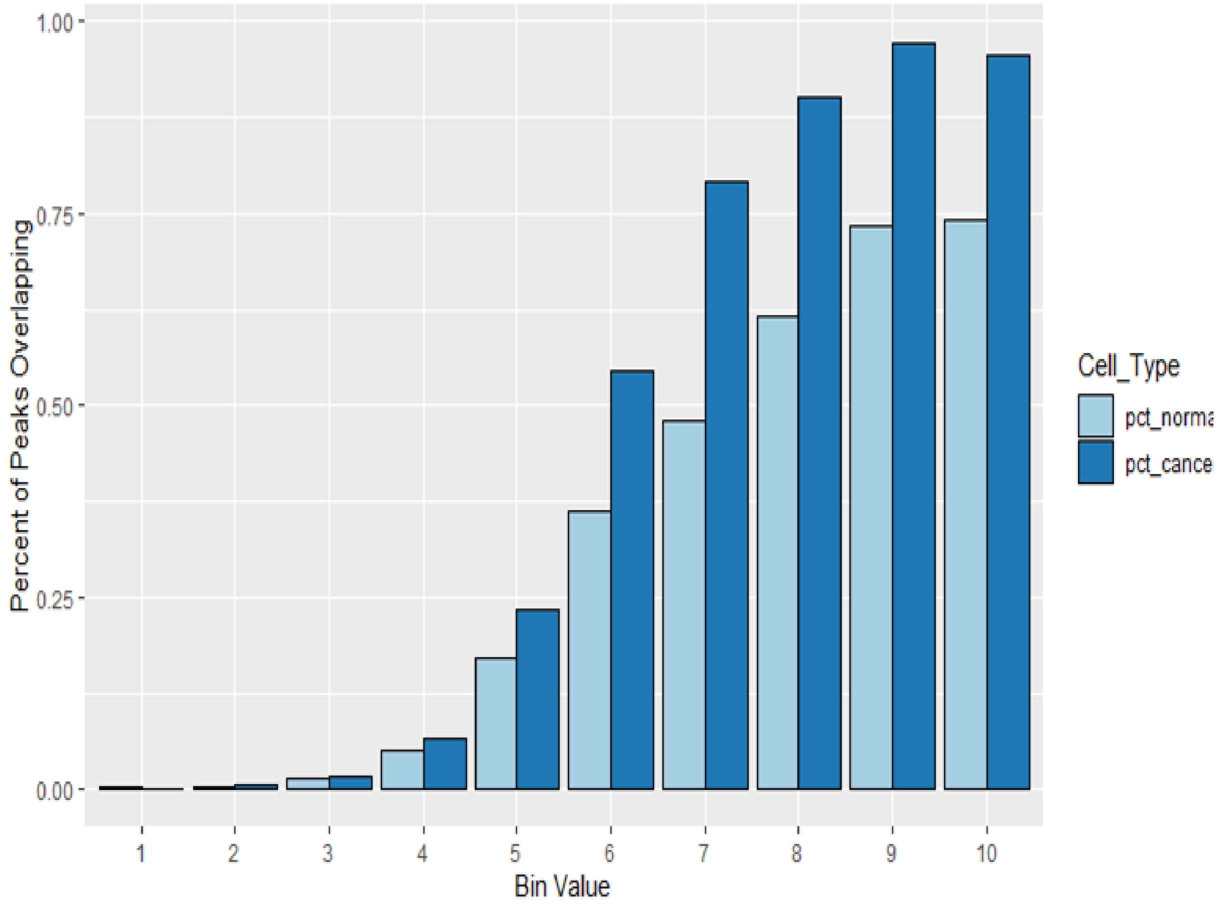

**Figure S11.** Comparison of continuous peaks with published consensus ChIP peaks. The fraction of intersection peaks over the total number of continuous peaks for each bin (Supplemental Figure S9) is plotted. The fraction values for normal cells (light blue) are compared with those for cancer cells (dark blue).

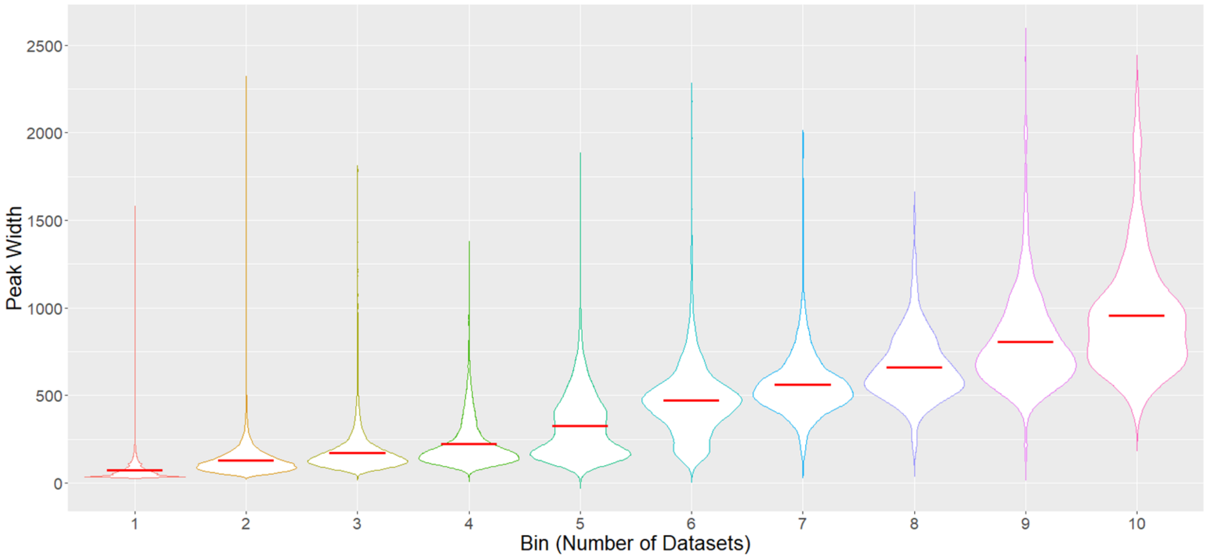

**Figure S12.** Peak width distribution. Peak widths across continuous peak bins are shown in the violin plot. No outlier peaks larger than 3000 bp width were included. All violins have been scaled to have maximum width values, and the red line indicates the mean value.

A.

| regulation | normal | cancer | chi-squared | |
|---|---|---|---|---|
| up | 287 | 208 | X-squared | 38.179 |
| down | 204 | 326 | df | 1 |
| | | | p-value | 6.46E-10 |

B.

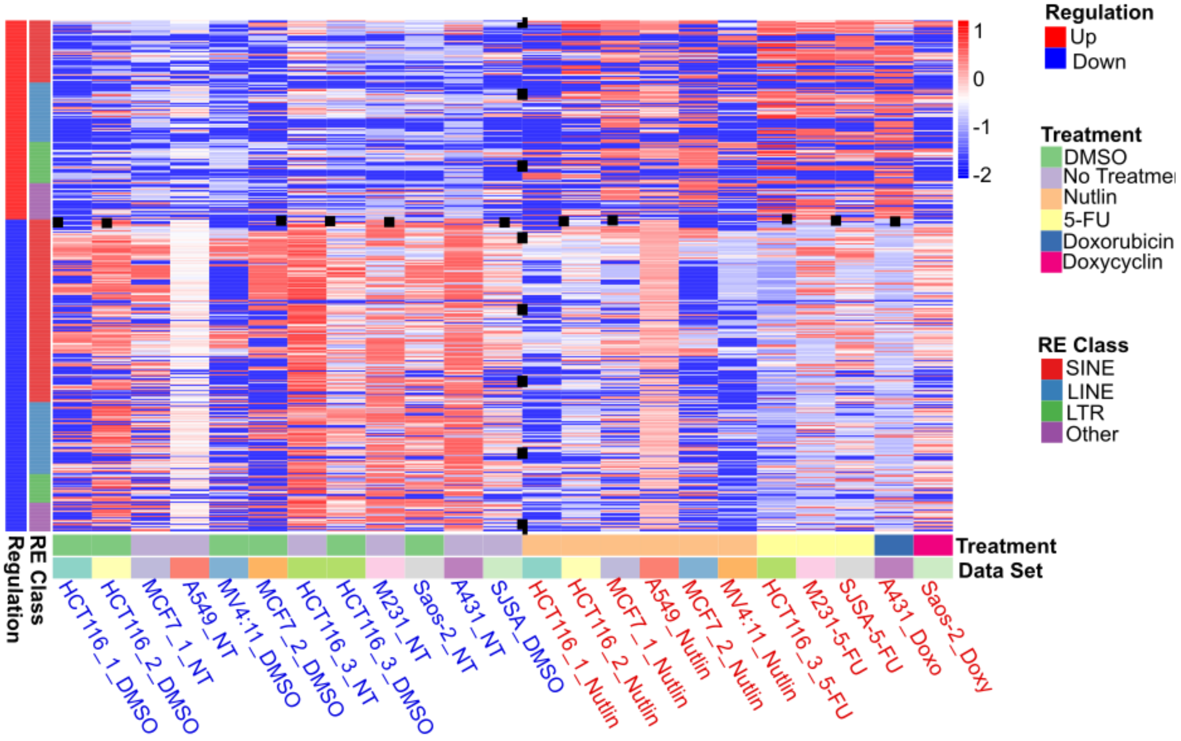

**Figure S13.** Comparison of p53-regulated TEs in cancer cells. (A) Summary of activated and repressed TEs. Statistical significance of the difference between normal and cancer cells is evaluated by Chi-squared test. (B) Heatmap for cancer cells. The heatmap is constructed to compare control (DMSO or non-treatment, blue) and treatment (red) samples, as well as TEs up- (red) and down-regulated (blue) by p53. Other notations are the same as those in Figure 5.

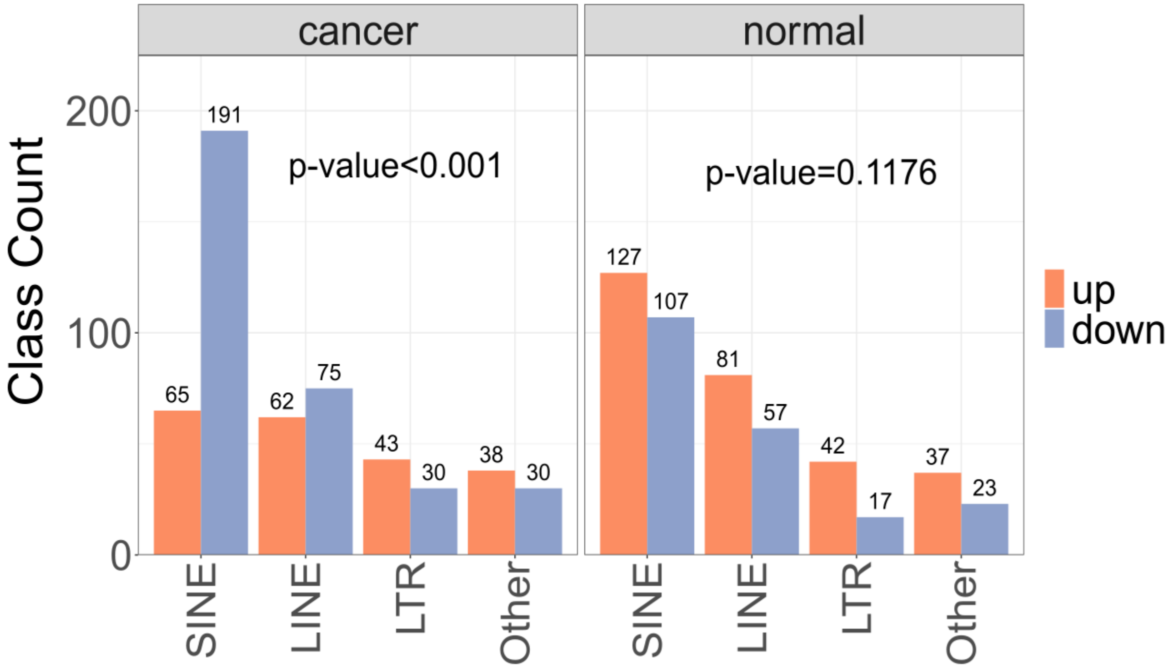

**Figure S14.** Repeat family analysis of p53-regulated TEs in normal and cancer cells. For each family, the number of up- (orange) and down-regulated (blue) TEs are shown. The statistical significance of the differences among repeat families is evaluated by Chi-squared tests.

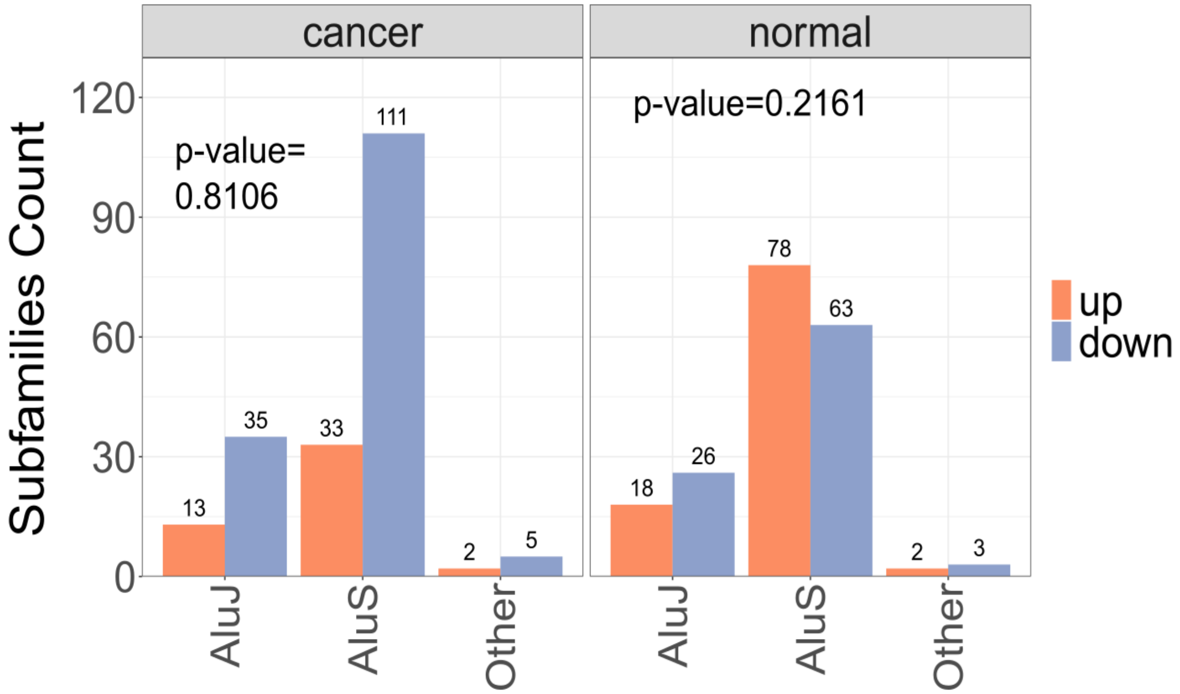

**Figure S15.** Alu subfamily analysis of p53-regulated TEs in normal and cancer cells. The notations are the same as Supplemental Figure S14.

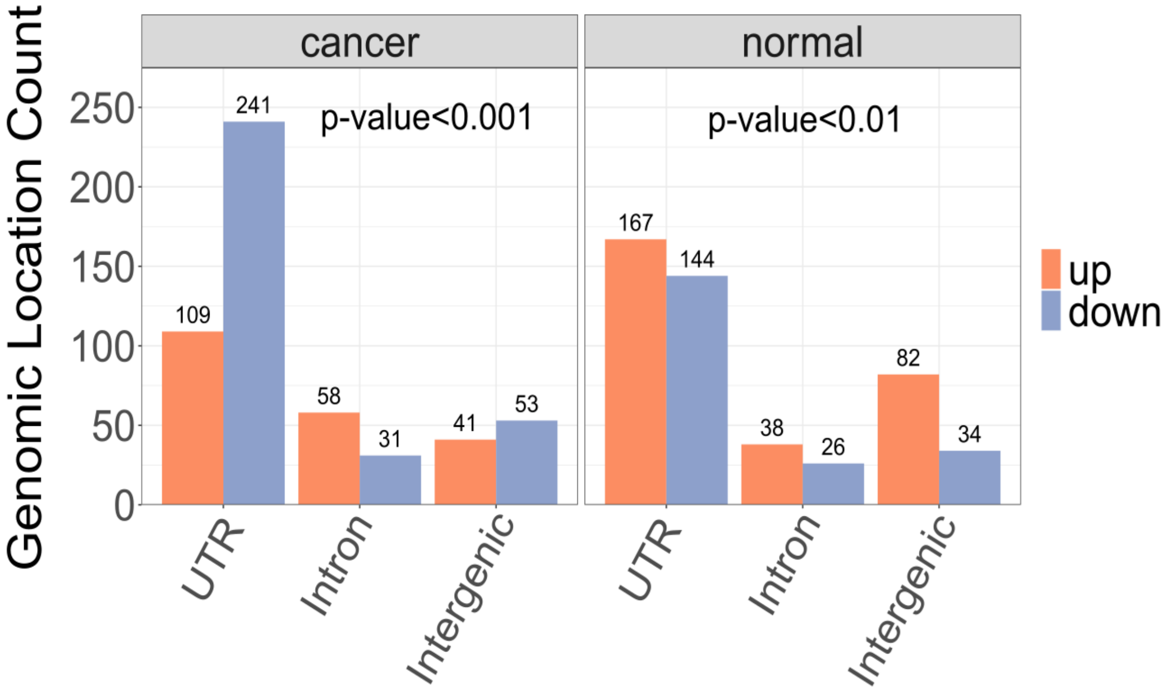

**Figure S16.** Genomic locations of p53-associated TEs. The numbers of the up- and down-regulated TEs in cancer (left) and normal (right) cells located in un-translated regions (UTR), introns and intergenic regions are compared. For each cell type, the statistical significance of the difference between up- and down-regulated TEs is evaluated by Chi-squared tests.
